# Induction of core symptoms of autism spectrum disorders by in vivo CRISPR/Cas9-based gene editing in the brain of adolescent rhesus monkeys

**DOI:** 10.1101/2020.08.03.233437

**Authors:** Shi-Hao Wu, Xiao Li, Dong-Dong Qin, Lin-Heng Zhang, Tian-Lin Cheng, Zhi-Fang Chen, Bin-Bin Nie, Xiao-Feng Ren, Jing Wu, Wen-Chao Wang, Ying-Zhou Hu, Yilin Gu, Long-Bao Lv, Yong Yin, Xin-Tian Hu, Zi-Long Qiu

**Affiliations:** Institute of Neuroscience, State Key Laboratory of Neuroscience, CAS Key Laboratory of Primate Neurobiology, Chinese Academy of Sciences, Shanghai, 200031, China; Kunming Institute of Zoology, Chinese Academy of Sciences, Kunming, Yunnan, 650223, China; Center for Excellence in Brain Science and Intelligence Technology, Chinese Academy of Sciences, Shanghai, 200031, China; Kunming Primate Research Center, Kunming Institute of Zoology, Chinese Academy of Sciences, Kunming, Yunnan, 650223, China; Kunming College of Life Science, University of the Chinese Academy of Sciences, Kunming, Yunnan, 650204, China; Beijing Engineering Research Center of Radiographic Techniques and Equipment, Institute of High Energy Physics, Chinese Academy of Sciences, Beijing 100049, China; School of Nuclear Science and Technology, University of Chinese Academy of Sciences, Beijing 100049, China; Department of Rehabilitation Medicine, Fourth Affiliated Hospital of Kunming Medical University, Kunming, Yunnan, 650021, China; Yunnan University of Traditional Chinese Medicine, 1076 Yuhua Road, Kunming, Yunnan, 650500, China; Academy for Engineering & Technology, Fudan University, Shanghai, 200433, China

**Keywords:** Autism spectrum disorders, Nonhuman primate model, disease model, Gene-editing

## Abstract

Although CRISPR/Cas9-mediated gene editing is widely applied to mimic human disorders, whether acute manipulation of disease-causing genes in the brain leads to behavioral abnormalities in non-human primates remains to be determined. Here we induced genetic mutations in MECP2, a critical gene linked to Rett syndrome (RTT) and autism spectrum disorders (ASDs), in the hippocampus (DG and CA1–4) of adolescent rhesus monkeys (Macaca mulatta) in vivo via adeno-associated virus (AAV)-delivered Staphylococcus aureus Cas9 with sgRNAs targeting MECP2. In comparison to monkeys injected with AAV-SaCas9 alone (n = 4), numerous autistic-like behavioral abnormalities were identified in the AAV-SaCas9-sgMECP2-injected monkeys (n = 7), including social interaction deficits, abnormal sleep patterns, insensitivity to aversive stimuli, abnormal hand motions and defective social reward behaviors. Furthermore, some aspects of ASDs and RTT, such as stereotypic behaviors, did not appear in the MECP2 gene-edited monkeys, suggesting that different brain areas likely contribute to distinct ASD symptoms. This study showed that acute manipulation of disease-causing genes via in vivo gene editing directly led to behavioral changes in adolescent primates, paving the way for the rapid generation of genetically engineered non-human primate models for neurobiological studies and therapeutic development.

## Introduction

Discovery of the clustered regularly interspaced short palindromic repeats (CRISPR)-associated protein (Cas9) system in bacteria and its optimization of genome editing have permitted efficient site-specific genetic manipulation in various mammalian cells and postmitotic tissues [1–4].

Gene editing has been successfully used in germline cells to create genetically engineered animals, including non-human primates, in order to mimic human brain disorders [5–9]. Since 2014, researchers have established genetically engineered non-human primates with CRISPR/Cas9 technology [10–12]. Moreover, various studies have identified the possibility of using CRISPR/Cas9 to generate non-human primate models for human disease [13–16].

In particular, adeno-associated virus (AAV)-delivered Staphylococcus aureus Cas9 (SaCas9) or Streptococcus pyogenes Cas9 (SpCas9) approaches can be applied to edit liver and muscle tissue genes in vivo with high efficiency [17–20], and viral vector-based gene editing is widely used in primate nervous system disease research, especially Parkinson’s disease [21–23]. However, it remains unknown whether direct in vivo gene editing in the adult brain of non-human primates can lead to ASD-related phenotypes, which would facilitate the modeling process and avoid the prolonged reproduction period required for primate species.

Genetic manipulation of MECP2, a critical gene in Rett syndrome (RTT) and autism spectrum disorders (ASDs), by lentiviral-based transgenic methods or transcription activator-like effector nuclease (TALEN)-based gene editing in non-human primates can result in autistic-like symptoms and faithfully mimic many aspects of MECP2-related disorders [9, 24–26]. However, their long reproduction cycles and inconvenience of international transfer have prevented the wide application of non-human primates as disease models. Thus, we investigated whether acute manipulation of the MECP2 gene via viral-delivered gene editing in adolescent rhesus macaque brains can lead to autism-related behavioral abnormalities, which should help facilitate disease modeling in non-human primates and address the role of MECP2 in particular brain regions, as well as assist in vivo neural circuitry research in non-human primates.

## Results

### Design of AAV-SaCas9 and sgRNA vectors and validation of efficiency of *in vivo* gene editing

Taking advantage of the small-sized Cas9 protein — *Staphylococcus aureus* Cas9 (SaCas9), we used AAV-mediated SaCas9 to delete the *MECP2* gene in the rhesus macaque brains *in vivo* and determine whether this leads to behavioral abnormalities [4]. To target the *MECP2* gene in the monkey genome, we synthesized two sgRNAs specifically targeting exon 4 in *MECP2* (Fig. 1A, B). The two sgRNAs (sgRNA-MECP2-a, b) were individually inserted into the AAV-hSyn-SaCas9-U6-sgRNA vector, with each harboring the human synapsin promoter to drive SaCas9 and U6 promoter to drive the sgRNA cassette. We used the AAV-hSyn-SaCas9-U6-sgRNA vector with sgRNA targeting the LacZ enzyme as the control vector.

**Figure 1.**
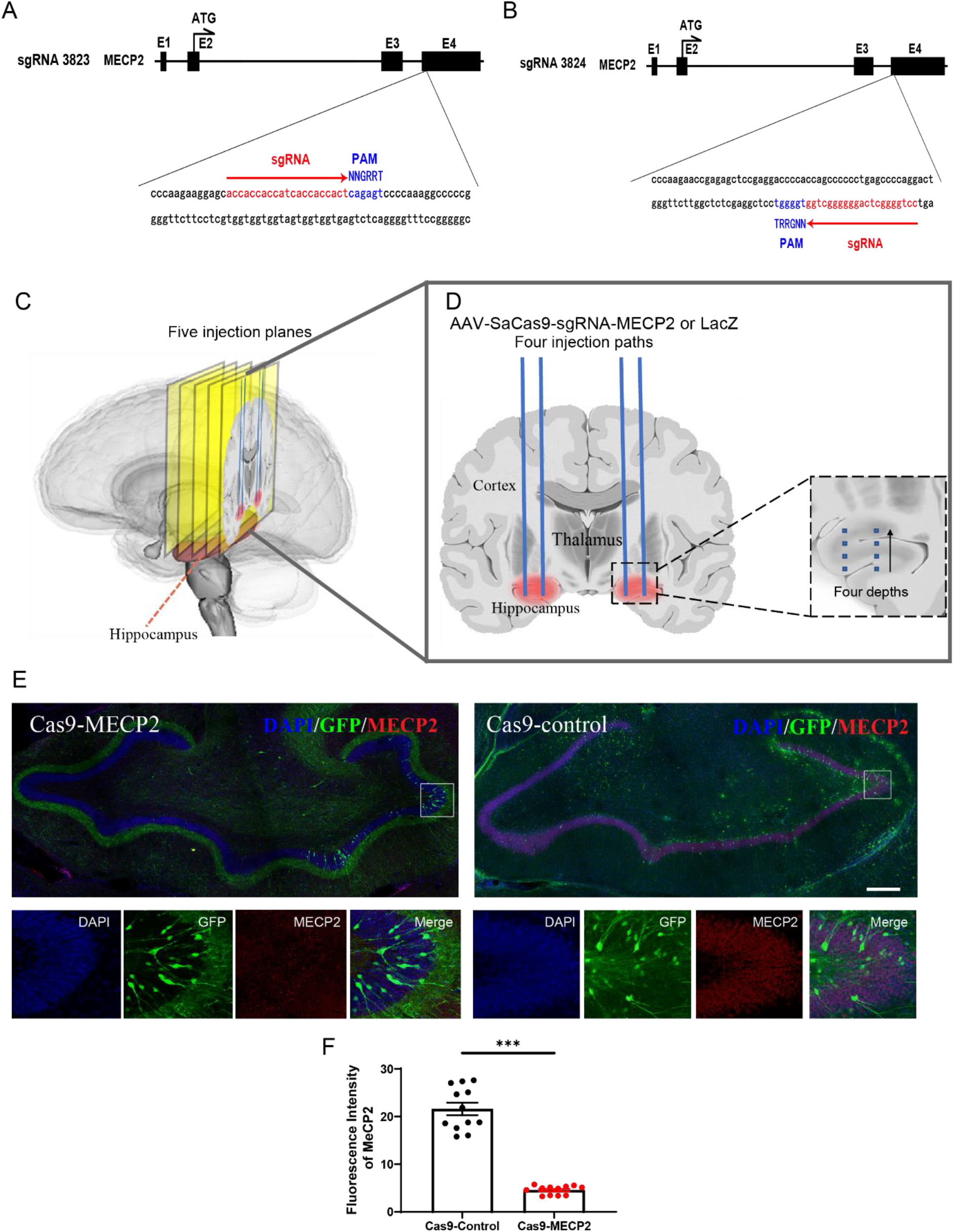
Design of sgRNAs for SaCas9, injection protocols, and validation *in vivo*. (**A-B**) Targeting sites of sgRNA for monkey *MECP2* gene. PAM sequences are in blue. sgRNA-MECP2-a was designed according to PAM sequence on plus stand and sgRNA-MECP2-b was designed according to PAM sequence on minus stand. (**C-D**) Schematic of injection strategy for monkey KO-2, location of hippocampus was determined by MRI images. Five injection planes were selected for viral injections from anterior to posterior to cover entire hippocampus. There were two injection paths in every injection plane on each side of the hippocampus. Injections were given at four different depth sites along each injection path. Details of injection strategy are provided in the Supplementary Materials. (**E**) Immunostaining of hippocampal sections of Cas9-MECP2 (KO-3) and Cas9-Control (Ctrl-2) monkeys for GFP (green), MECP2 (red), and DAPI (blue). Scale bar, 500 μm. (**F**) Arbitrary units for fluorescence intensity of MECP2 protein signals in DAPI-positive cells detected by anti-MECP2 antibody were quantified (n = 12, from two monkeys of each group, Cas9-Control: Ctrl-1 and Ctrl-2, Cas9-MECP2: KO-3 and KO-7). Fluorescence signal of MECP2 proteins was reduced in Cas9-MECP2 monkeys (Mann-Whitney test, U = 0, *** *p* < 0.0001).

To ensure complete deletion of *MECP2*, we injected two AAV-hSyn-SaCas9-U6-sgMECP2-a/b vectors and AAV-hSyn-GFP (green fluorescent protein indicating injection site) into the Cas9-MECP2 monkey group (n = 7, age: 2–3 years). We injected AAV-hSyn-SaCas9-U6-sgLacZ and AAV-hSyn-GFP into the Cas9-Control monkey group (n = 4, age: 2–3 years) (Table S1A–I).

The hippocampus is recognized as a critical region for social memory [27, 28]. Thus, we examined whether deletion of *MECP2* in the monkey hippocampus leads to autistic-like phenotypes. To localize the hippocampus precisely in the monkey brain, a magnetic resonance imaging (MRI)-based deep brain structure localization procedure (with an error < 1.0 mm) was applied (Fig. S1). After localization, we injected the AAVs into ~40 sites on each side of the hippocampus (Fig. 1C, D, see Table S2 for injection coordinates and injected virus volumes, and figure legends and Methods for technical details) [29, 30].

To confirm the efficiency of gene editing, we amplified the sgRNA-targeting genomic regions from monkey hippocampal tissues via polymerase chain reaction (PCR). We identified critical genetic mutations in these regions, including stop-codons, thus showing the effectiveness of SaCas9-mediated gene editing (Fig. S2). To further determine whether the MECP2 protein level was altered by gene editing, we performed immunohistochemical experiments on sections of the monkey hippocampus. Results showed that the MECP2 protein level decreased significantly in the hippocampus of the Cas9-MECP2 monkeys compared with that of the Cas9-Control monkeys (Fig. 1E-F). This indicated that the CRISPR/Cas9 system effectively reduced the expression of endogenous MECP2 in the monkey brain via gene editing. To rule out the possibility of SaCas9-mediated off-target genome editing, we identified potential genomic off-target sites in the hippocampal tissue of injected monkeys. However, the top three off-target site candidates of each sgRNA showed no signs of off-site genome editing in protein-coding regions (Fig. S3).

To determine whether viral injections in the monkey brain cause unexpected neuroinflammation effects and inflammation-mediated destruction of transduced cells, immunohistochemical experiments were performed on the monkey hippocampal slides. We found that both astrocyte and microglial cells exhibited negative activation signals, e.g., numbers of GFAP- and Iba1-positive cells as well as process lengths and endpoints of Iba1-positive cells (Fig. S4A-F).

### Examination of social behaviors within a community

According to the Diagnostic and Statistical Manual of Mental Disorders (5th Edition, DSM-V) and clinical diagnostic criteria for RTT and ASDs [31–33], we designed several behavioral assays to analyze the core behavioral phenotypes of monkeys after viral injection for 3–6 months (Table S1A). We first compared the behavioral phenotypes for individual monkeys before and after viral injection. As the rhesus monkeys were not inbred strains, comparison of behaviors with or without manipulation of individual monkeys can serve as a reliable parameter. We also examined the role of *MECP2* in these behaviors by comparing the gene-edited group (Cas9-MECP2) with the control group (Cas9-Control). Abnormality in social interactions and communication are hallmarks of ASDs. To observe the natural behaviors of monkeys within a social community, we established several social groups (each consisting of 6–8 age-matched male monkeys, including one or two gene-edited monkeys). All behaviors were video-recorded and then analyzed off-line to examine the various behaviors of individual monkeys before and after gene editing, as well as between the Cas9-MECP2 and Cas9-Control groups [9].

Firstly, the duration of active social contact, defined as initiating play, sharing toys, grooming others, and sitting together [9], was significantly decreased after the Cas9-MECP2 virus injection (monkey number, n = 7) compared to baseline prior to viral injection (Movie S1, S2, S3), but was not observed in the Cas9-Control group (monkey number, n = 3) (Fig. 2A-C). After gene editing, the Cas9-MECP2 group also exhibited significantly less active social contact compared with the Cas9-Control group (Fig. 2D). The frequency of active social contact was not altered with injection of either Cas9-MECP2 or Cas9-Control virus (Fig. 2E, F). The frequency of active social contact in the Cas9-MECP2 group displayed a decreasing (but non-significant) trend compared with the Cas9-Control group (Fig. 2G, H). The significant difference in social interaction before and after injection of the Cas9-MECP2 virus suggests that *MECP2* in the hippocampus plays a critical role in regulating social interaction in monkeys.

**Figure 2.**
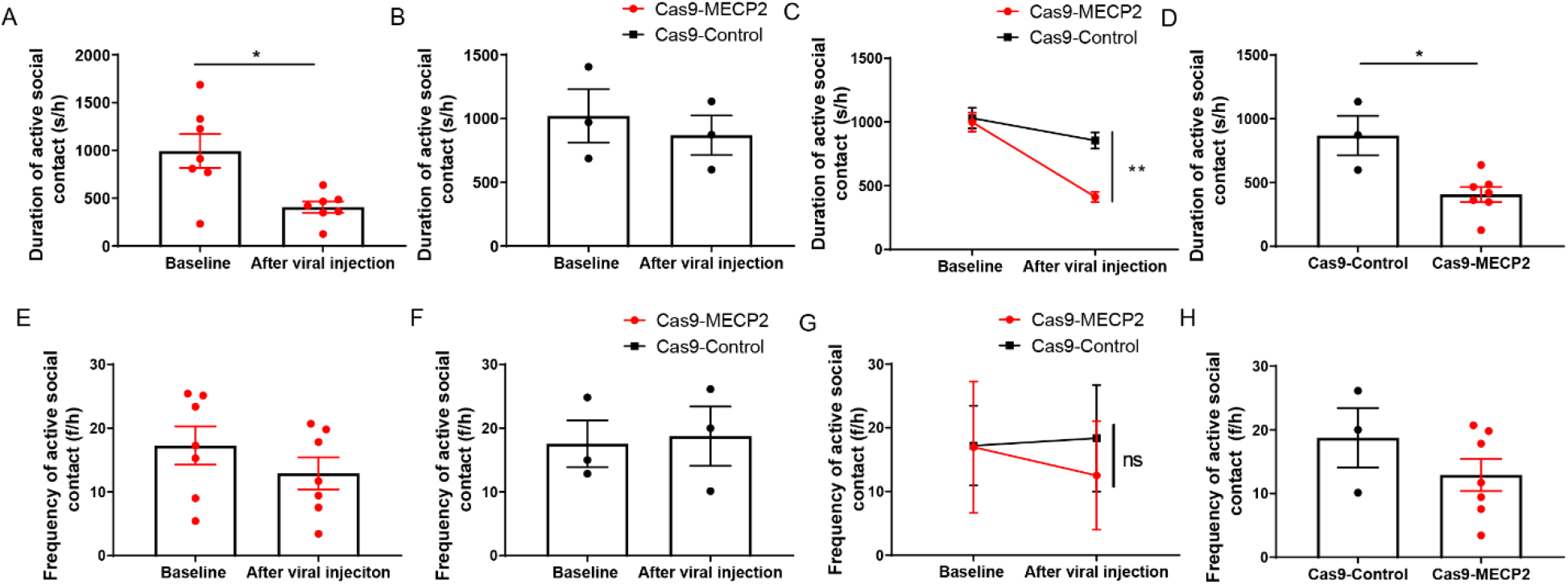
Monkeys with acute *MECP2* gene editing exhibited abnormal social behaviors. (**A**) Duration of social contact in Cas9-MECP2 group (Mann-Whitney test, U = 6, * *p* = 0.0175, n = 7, from seven monkeys, average of 7 d of repeated measurements per monkey) and (**B**) Cas9-Control group before and after injection (n = 3, from three monkeys, average of 7 d of repeated measurements per monkey). (**C**) Changes in active social contact after injection were significantly different between Cas9-MECP2 and Cas9-Control groups (repeated-measures ANOVA, F = 8.245, ** *p* = 0.005, η_p_^2^ = 0.111). (**D**) Cas9-MECP2 group (n = 7) showed significantly shorter duration of active social contact compared to Cas9-Control group (n = 3) after injection (Mann-Whitney test, U = 1, * *p* = 0.0333). (**E**) Frequency of social contact in Cas9-MECP2 group (n = 7, from seven monkeys, average of 7 d of repeated measurements per monkey) and Cas9-Control group **(F)** (n = 3, from three monkeys, average of 7 d of repeated measurements per monkey). (**G**) No significant differences in frequency were found between groups (repeated-measures ANOVA, p = 0.115). (**H**) Cas9-MECP2 group (n = 7) showed similar frequency of active social contact as Cas9-Control group (n = 3) after injection. Each spot represents average duration time of social contact (7 d) for individual monkeys during daily observation periods. Data are means ± SEM, data within 95% confidence interval were included in statistical analysis.

### Altered aggressive-like behaviors in monkeys following *MECP2* gene editing

It is well documented that ASD individuals in human communities are often at a higher risk of being bullied [34]. We reasoned that, due to altered social behaviors, the *MECP2* gene-edited monkeys may hold inferior social positions and, as a result, display less aggressive behaviors. Thus, we determined whether aggressive and submissive behaviors were altered in gene-edited monkeys in the social community described above. We found that active attack behaviors, such as biting, slapping, grabbing, and stare threatening, were significantly decreased in monkeys injected with the Cas9-MECP2 virus, which was not found in the Cas9-Control group (Fig. 3A, B, C). The Cas9-MECP2 group exhibited significantly less active aggressive behaviors compared with the Cas9-Control group after viral injection (Fig. 3D).

**Figure 3.**
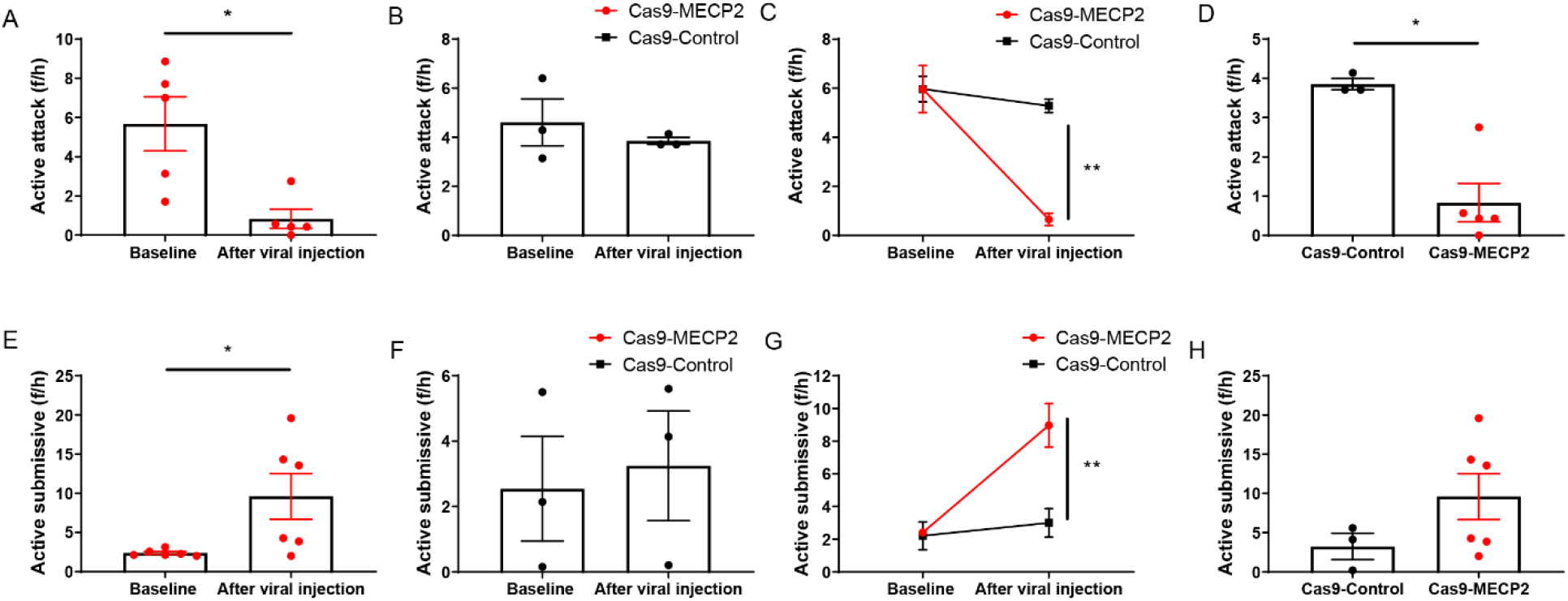
Monkeys with acute *MECP2* gene editing exhibited abnormal aggressive behaviors. (**A**) Active attacks decreased in Cas9-MECP2 group (n = 5, from five monkeys, average of 7 d of repeated measurements per monkey) after injection (Mann-Whitney test, U = 1, * *p* = 0.0159), (**B**) but not in Cas9-Control group (n = 3, from three monkeys, average of 7 d of repeated measurements per monkey). (**C**) Changes in active attacks after injection were significantly different between Cas9-MECP2 and Cas9-Control groups (repeated-measures ANOVA, F = 12.232, ** *p* = 0.0010, η_p_^2^ = 0.200). (**D**) Cas9-MECP2 group (n = 5) showed significantly lower active attacks than Cas9-Control group (n = 3) after injection (Mann-Whitney test, U = 0, * *p* = 0.0357). (**E**) Submissive behaviors increased in Cas9-MECP2 group after injection (n = 6, from six monkeys, average of 7 d of repeated measurements per monkey, Mann-Whitney test, U = 5.5, * *p* = 0.0476), (**F**) but not in Cas9-Control group (n = 3, from three monkeys, average of 7 d of repeated measurements per monkey). (**G**) Changes in submissive behaviors were significantly different between Cas9-MECP2 and Cas9-Control groups (repeated-measures ANOVA, F = 8.022, ** *p* = 0.006, η_p_^2^ = 0.125). (**H**) Cas9-MECP2 group (n = 6) displayed more submissive behaviors than Cas9-Control group (n = 3) after injections. Each dot indicates average time (7 d) of social activity for individual monkeys during daily observation period. Data are means ± SEM, data within 95% confidence interval were included in statistical analysis.

Consequently, submissive behaviors, such as screaming threat, crouching, fleeing, lip smacking, grimacing, submissive presenting, or escaping, increased remarkably after injection of the Cas9-MECP2 virus compared to injection of the Cas9-Control virus (Fig. 3E, F, G, H). This evidence indicates that Cas9-MECP2-mediated gene editing significantly altered social interactions, strongly suggesting that manipulation of the *MECP2* gene is sufficient to modulate social behaviors in adolescent monkeys.

### Abnormal circadian activity and sleep patterns in monkeys following *MECP2* manipulations

Sleep problems are a prominent feature of ASDs and RTT [32, 35–37]. To address whether sleep patterns can be altered by *in vivo MECP2* gene editing, we used collar-like monitors (Actical physical activity monitors, Respironics, Pennsylvania, USA) belt-fixed onto the necks of the monkeys for 7–9 d (according to the battery life) to record 24-h activity (Table S1B). After offline analysis, we found that monkeys injected with the Cas9-MECP2 virus (n = 3) showed decreased locomotor activity during the day compared to monkeys injected with the Cas9-Conrol virus (n = 3) (Fig. 4A, B, C, D). To further measure the sleep patterns of the gene-edited monkeys, activity data during the night (19:00 pm–07:00 am) were classified into awake and sleep phases, with the sleep phase further divided into relaxed (no body movements during sleep) and transitional (1–2 movements within 1 min) sleep phases (see Methods for definitions).

**Figure 4.**
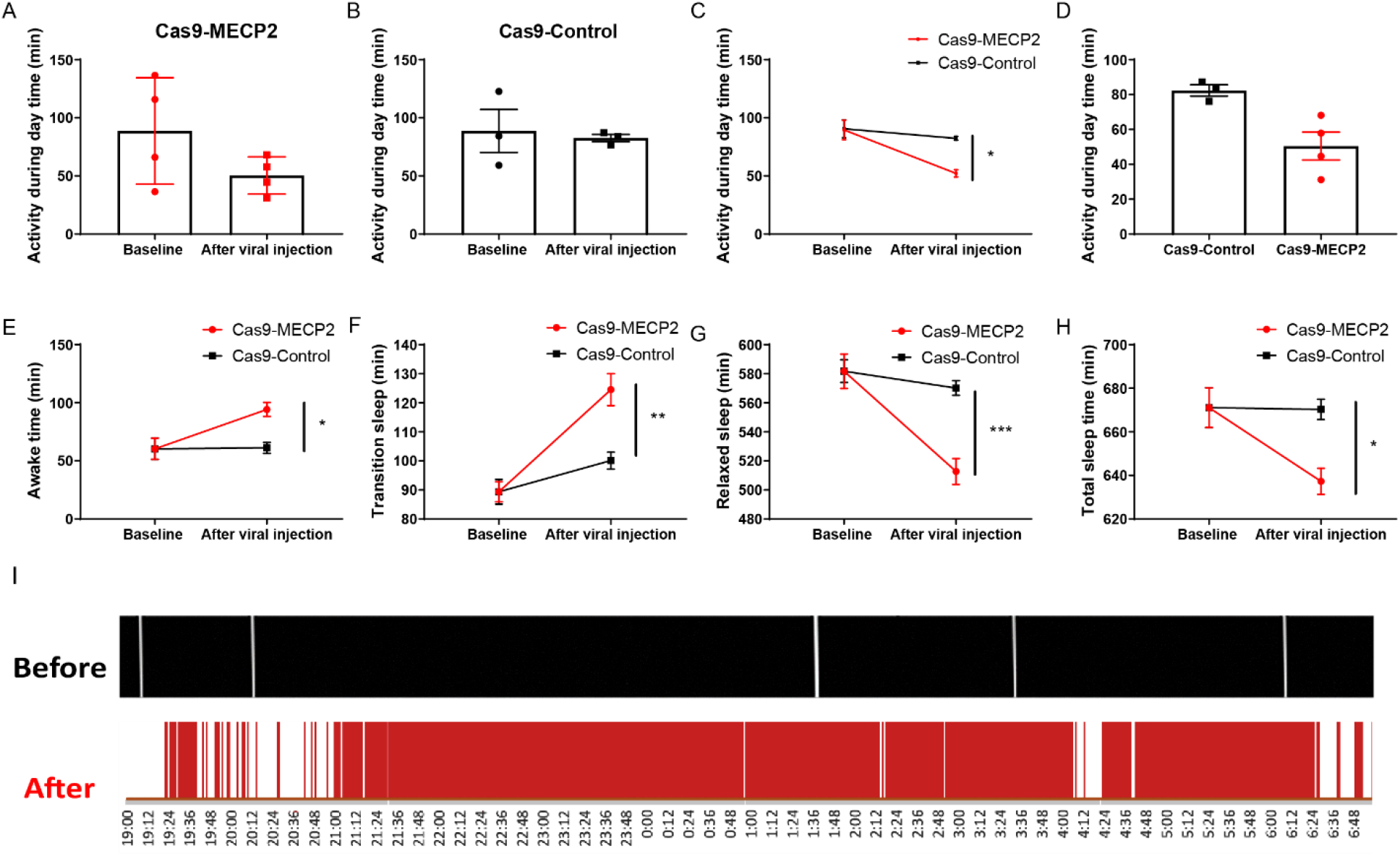
Hypoactivity and abnormal sleep patterns in acute *MECP2* gene-edited monkeys. (**A**) Cas9-MECP2 (n = 4, from four monkeys, average of 7 d of repeated measurements per monkey) and (**B**) Cas9-Control groups (n = 3, from three monkeys, average of 7 d of repeated measurements per monkey) exhibited no significant changes in daytime activity. (**C**) Changes in daytime activity after viral injection were significantly different between Cas9-MECP2 and Cas9-Control groups (repeated-measures ANOVA, F = 6.782; * *p* = 0.013; η_p_^2^ = 0.136). (**D**) After viral injection, Cas9-MECP2 group showed shorter daytime activity compared with Cas9-Control group. (**E**) Cas9-MECP2 group showed significantly increased awake time duration after injection compared with control group (repeated-measures ANOVA, F = 6.782; * *p* = 0.012, η_p_^2^ = 0.126; Cas9-MECP2 group, n = 28, from 7 d of measurements of four monkeys; Cas9-Control group, n = 21 from 7 d of measurements of three monkeys). (**F**) Cas9-MECP2 group showed significantly increased duration of transitional sleep compared with control group (repeated-measures ANOVA, F = 10.6, ** *p* = 0.002, η_p_^2^ = 0.184; Cas9-MECP2 group, n = 28, from 7 d of measurements of four monkeys; Cas9-Control group, n = 21 from 7 d of measurements of three monkeys). (**G**) Cas9-MECP2 group exhibited significantly decreased duration of relaxed sleep compared with control group (repeated-measures ANOVA, F = 14.04; *** *p* < 0.0001, η_p_^2^ = 0.23; Cas9-MECP2 group, n = 28, from 7 d of measurements of four monkeys; Cas9-Control group, n = 21, from 7 d of measurements of three monkeys). (**H**) Cas9-MECP2 group exhibited significantly decreased duration of total sleep compared with control group (repeated-measures ANOVA, F = 6.782, * *p* = 0.012; η_p_^2^ = 0.126; Cas9-MECP2 group, n = 28, from 7 d of measurements of four monkeys; Cas9-Control group, n = 21 from 7 d of measurements of three monkeys). (**I**) Two examples of sleep patterns (12 h from 19:00 pm to 07:00 am) from same monkey (KO-3), illustrating fragmented sleep after gene editing. Each dot (in **A, B, D**) indicates average of 7 d activity of individual monkeys during 12-h daily observation period (07:00 am–19:00 pm, 7 d). Data are means ± SEM.

We found that durations of the awake and transitional sleep phases were significantly increased in monkeys (n = 4) injected with the Cas9-MECP2 virus compared to monkeys injected with the Cas9-Control virus (Fig. 4E, F). Consequently, the relaxed sleep period and overall sleep time decreased in the Cas9-MECP2 virus-injected monkeys, but not in monkeys injected with the Cas9-Control virus (Fig. 4G, 4H). Thus, sleep patterns were significantly fragmented in monkeys (e.g., #KO-3) injected with the Cas9-MECP2 virus (Fig. 4I). These abnormalities in sleep and activity rhythms are similar to those observed in autistic patients [32, 37, 38] and to the phenotypes identified in TALEN-edited *MECP2*-mutant cynomolgus monkeys [9].

### Defective social reward and communication behaviors in monkeys with *MECP2* deletions

One prevalent hypothesis for the development of autistic-like behavioral abnormalities is that ASD patients do not gain sufficient reward from social interaction, thereby leading to diminished motivation for seeking or maintaining social contact [39]. Thus, we investigated whether social reward is compromised by *MECP2* editing in the hippocampus using a monkey conditioned place preference (CPP) paradigm [40] (Fig. S5, Table S1C). Briefly, monkeys underwent two conditioning steps (i.e., social isolation and social interaction), followed by the choice to enter either the isolation chamber (assigned based on previous color preference) or social chamber (where a female partner was present during previous social conditioning). After social conditioning, both groups of monkeys spent more time in the social cages than before (Fig. 5A, B). However, the Cas9-Control monkeys (n = 3) spent more time in the social cages than the Cas9-MECP2 monkeys (n = 3), though without statistical significance (Fig. 5C, Movie S4, S5). These results suggest that 3 d of social conditioning had a mild effect on the Cas9-MECP2 monkeys, but an obvious effect on the Cas9-Control monkeys, supporting our speculation that deletion of *MECP2* in the hippocampus compromises the social reward system, which may further aggravate social interaction abnormalities in monkeys.

**Figure 5.**
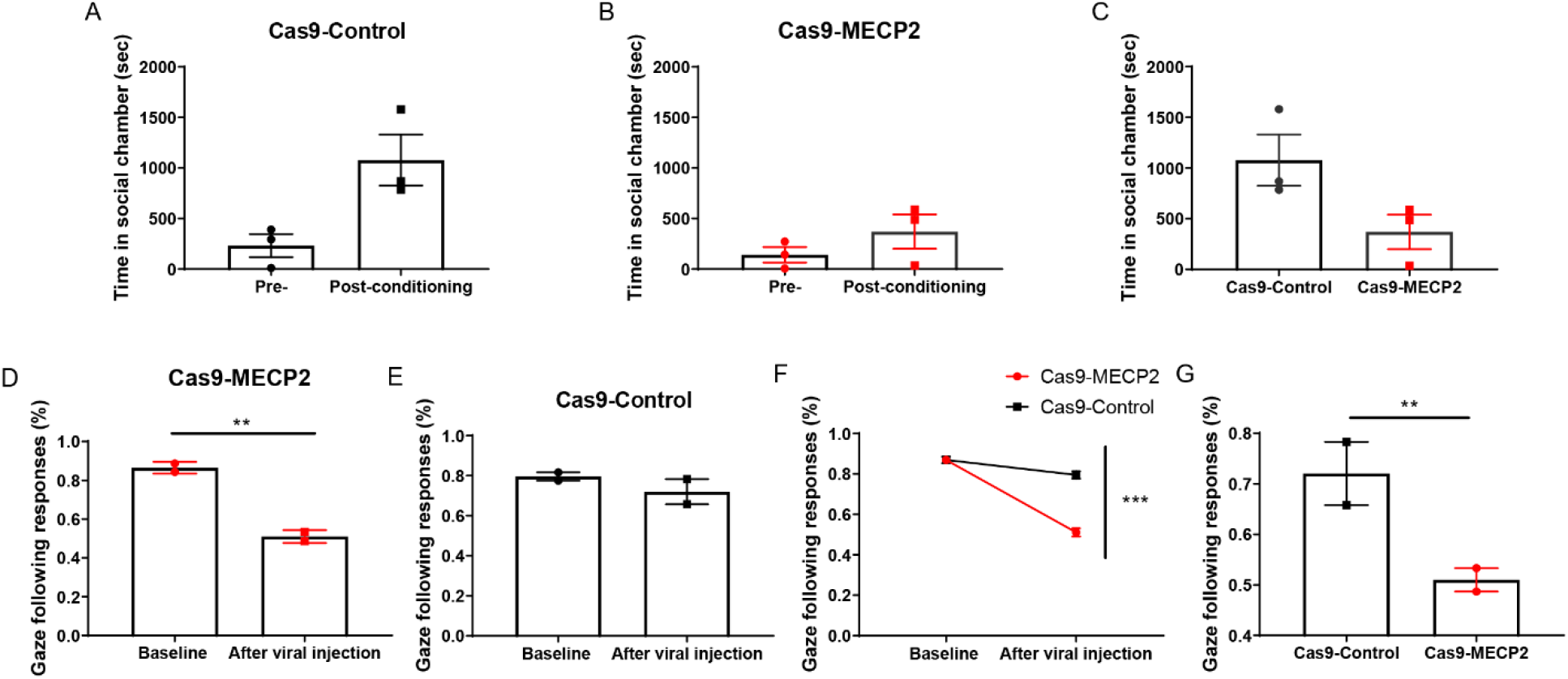
Monkeys with *MECP2* gene editing exhibited deficits in social reward and gaze following. After isolation and social conditioning, (**A**) Cas9-Control (n = 3 monkeys) and (**B**) Cas9-MECP2 groups (n = 3 monkeys) exhibited no significant changes in duration in social chamber. **(C)** Compared to Cas9-MECP2 group, Cas9-Control group spent more time in social chamber, revealing possible deficits in social reward function. (**D**) Cas9-MECP2 group showed significantly decreased gaze-following after virus injection (Mann-Whitney test, U = 0, ** *p* = 0.0022, n = 6, from 3 d of measurements of two monkeys). (**E**) Cas9-Control group exhibited no significant changes in gaze-following responses (n = 6, from 3 d of measurements of two monkeys). (**F**) Changes in gaze-following rate were significantly different between Cas9-MECP2 and Cas9-Control groups (repeated-measures ANOVA, F = 21.114, *** *p* = 0.001, η_p_^2^ = 0.679). (**G**) Cas9-MECP2 group showed significantly lower gaze-following rate than Cas9-Control group after viral injection (Mann-Whitney test, U = 1, ** *p* = 0.0022). (**A, B**) Each dot indicates duration of individual monkeys in social chamber during 30-min testing period, and (**D, E, G**) each dot indicates average percentage of gaze-following of individual monkeys during 3-d test period. Data are means ± SEM.

To further investigate social communication abilities, we observed the gaze-following behavior of monkeys elicited by tracking the gaze of human experimenters [41]. Gaze-following is defined as the ability of an individual to follow the direction of another’s gaze. This ability is a critical process through which social animals acquire necessary information on the emotional and mental states of social partners. The age at which a human infant can follow the gaze of a caregiver ranges from 6 to 18 months, with monkeys also developing this ability during infancy [41, 42]. However, children with ASD usually manifest developmental abnormality in gaze-following ability as a form of social communication defect [42, 43]. Our results revealed that gaze-following responses in monkeys (n = 2) decreased significantly following injection with the Cas9-MECP2 virus, but this was not observed in the Cas9-Control monkeys (n = 2) (Fig. 5D, E, F, Movie S6, S7, Table S1D). The Cas9-MECP2 group also showed significantly lower gaze-following than the Cas9-Control group after virus injection (Cas9-MECP2 monkeys: n = 2, Cas9-Control monkeys: n = 2, Fig. 5G, Movie S6, S7, Table S1D). The differences in gaze-following were not caused by different interactions between the monkey and trainer as monkeys injected with either the Cas9-MECP2 or Cas9-Control virus spent similar durations facing and looking at the trainer (Fig. S6A, B).

### Insensitivity for aversive sensory stimuli and lack of stereotypic behaviors in monkeys with *MECP2* manipulations

A common feature of ASD patients is their abnormal response to sensory stimuli (i.e., hyper- or hypo-sensitivity), which can be a precipitating factor for deficits in social skills [44, 45]. To determine whether *MECP2* gene editing in the hippocampus leads to defects in sensory responses, we applied an active avoidance test to examine the auditory sensory responses of the monkeys [9]. The tested monkeys were placed in a shuttle box with two connected chambers, each equipped with a high-pitch loudspeaker. High decibel white noise (116 dB) was played in the chamber housing the tested monkey, which would normally trigger an escape response. Compared with the Cas9-Control monkeys (n = 2), the Cas9-MECP2 monkeys (n = 2) became insensitive to noise after virus injection (Fig. 6A, B, C, Movie S8, S9, Table S1E). Specifically, the Cas9-MECP2 group exhibited much longer escape time than the Cas9-Control group (Cas9-MECP2 monkeys: n = 2, Cas9-Control monkeys: n = 2, Fig. 6D), suggesting that *MECP2* deletion in the hippocampus led to insensitivity to aversive sensory stimuli, thereby mimicking the sensory abnormalities of ASD patients. Interestingly, this result is similar to the abnormalities observed in whole-body TALEN-edited *MECP2*-mutant monkeys under the same procedure, suggesting that auditory sensory response abnormalities may be mediated by hippocampus-related neural circuitry [9].

**Figure 6.**
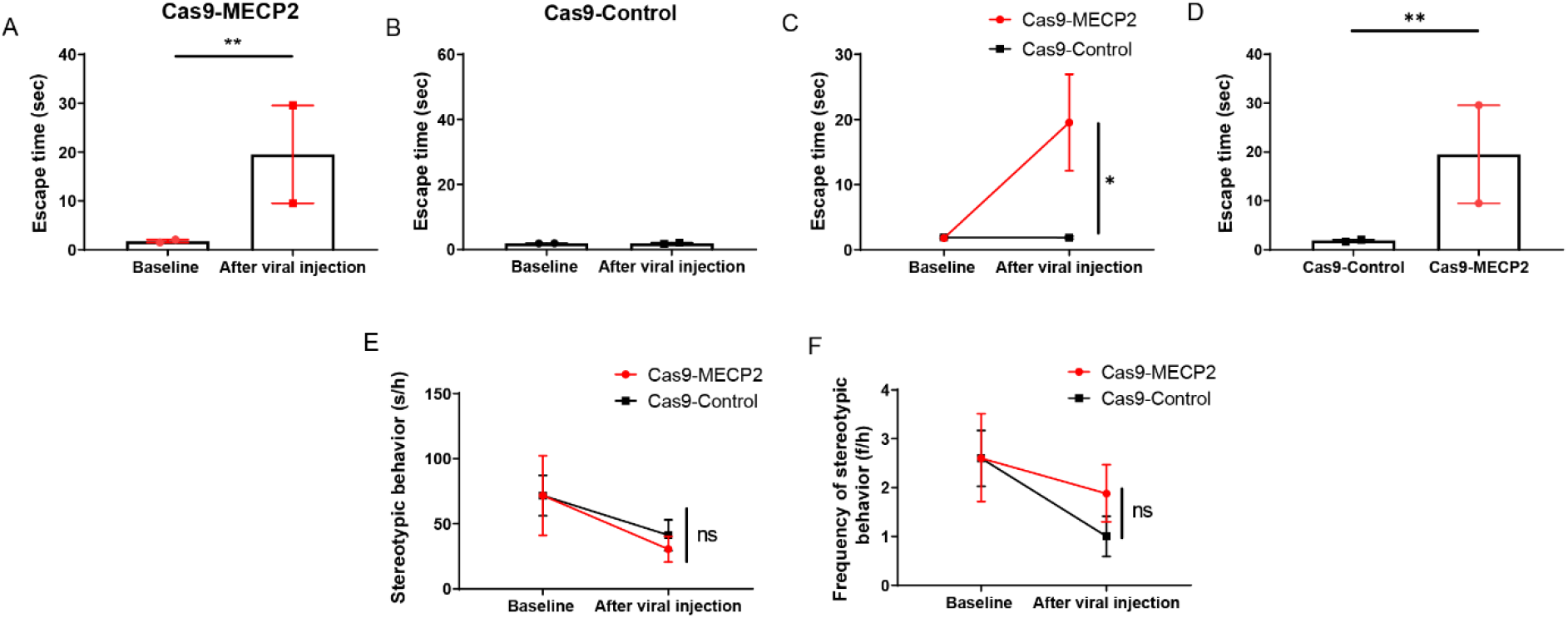
Monkeys with *MECP2* gene editing exhibited abnormal responses to aversive sensory stimuli and unchanged repetitive behaviors. (**A**) After injection, Cas9-MECP2 group showed significantly increased escape latency from noise chamber in active avoidance test (Mann-Whitney test, U = 0, ** *p* = 0.0022, n = 6, from 3 d of measurements of two monkeys). (**B**) Cas9-Control group exhibited no change in escape latency (n = 6, from 3 d of measurements of two monkeys). (**C**) Changes in both groups were significantly different (repeated-measures ANOVA, F = 5.644, * *p* = 0.039, η_p_^2^ = 0.361; Cas9-MECP2, n = 6; Cas9-Control, n = 6). (**D**) Cas9-MECP2 group exhibited significantly longer escape latency than Cas9-Control group after injection (Mann-Whitney test, U = 0, ** *p* = 0.0022). Duration (**E**) and frequency (**F**) of stereotypic behaviors exhibited no significant changes in Cas9-MECP2 and Cas9-Control groups (repeated-measures ANOVA, E: *p* = 0.833, F: *p* = 0.29). (**A, B, D**) Each dot indicates average escape latency of individual monkeys during 3-d observation period. Data are means ± SEM.

Stereotypic and repetitive behaviors are core symptoms in ASDs and RTT[33]. Importantly, animal models based on *MECP2* mutation or duplication, including non-human primate models, exhibit various stereotypic behaviors [9, 26]. In the present study, however, stereotypic behaviors were not observed in monkeys injected with either the Cas9-MECP2 or Cas9-Control virus (Fig. 6E, F, Table S1A). These data suggest that manipulation of *MECP2* in the hippocampus specifically affects some aspects of social behavior and stereotypic behaviors may be regulated by the thalamic reticular system, as suggested previously [46]

### Abnormal hand motions in monkeys with *MECP2* manipulations

Severe impairment in motor functions, especially abnormal hand motions, are primary components of Rett syndrome[31]. To examine whether *MECP2* gene editing in the brain causes motor disabilities in monkeys, we measured their physical mobility by the gait speed test (Fig. S7A) [47] and fine hand coordination by a modified brinkman board test (Fig. S7B) [48]. No significant differences between the gene-edited and control groups were found in gait speed test, but there was significant difference in brinkman board test. Compare to control monkeys, gene-edited monkeys take longer times to retrieve a peanut kernel from designed slots, which showed the abnormal hand motions in gene-editing monkeys (Fig. S7A, B, Table S1F, Table S1G). Therefore, we concluded that down-regulation of the *MECP2* gene in the hippocampus will not cause postural dyskinesia, but will induce fine hand coordination disabilities in the rhesus monkeys.

### *MECP2* edited monkeys may have learning and memory retrieval problems

As Rett syndrome patients also exhibit severe impairment of cognitive functions, we performed two further behavioral tests to examine whether deletion of *MECP2* affects cognition and memory in monkeys. A reversal learning task was used to assess learning and cognitive flexibility and a modified pattern-recognition task was used to test memory [49, 50]. Although no statistical differences were found between the Cas9-MECP2 (n = 3) and Cas9-Control monkeys (n = 2) in these two tasks (Fig S8A, B, S9A–C, Table S1H–I), the Cas9-MECP2 monkeys needed more trials to complete the learning phase and achieved a lower correct rate in the memory retrieval phase. This evidence suggests that Cas9-MECP2 monkeys may have learning and memory retrieval problems. Regrettably, due to the limitation of monkey numbers, the observed decline in cognition and memory in the Cas9-MECP2 monkeys did not achieve a statistical conclusion. Thus, whether down-regulation of the *MECP2* gene in the hippocampus causes impairment of cognition and memory requires further study.

### *MECP2* edited monkeys exhibited autistic-like phenotypes two years after gene-editing

In order to examine the stability of the phenotype, longitudinal analysis was performed in the rest of gene-editing and control monkeys. We analyzed social behaviors, light/dark activity, and sleep patterns for 5 d on monkeys 2–3 years after gene-editing surgery (Table S4). As shown in the following figures (Fig S10), we found that Cas9-MECP2 monkeys consistently exhibited social deficit phenotypes, measured by duration of social interaction time and time monkeys spent alone in comparison with Cas9-Control monkeys (Fig S10A). In the light/dark activity and sleep pattern tests, we found that Cas9-MECP2 monkeys still showed increased transitional sleep and decreased relaxed sleep compared to Cas9-Control monkeys. The day-time activity of Cas9-MECP2 monkeys also decreased compared to Cas-Control monkeys, consistent with the phenotypes (Fig S10B). Therefore, we concluded that *in vivo* gene editing in monkey brains had lasting effects regarding social behaviors, as well as circadian activities.

## Discussion

We showed that AAV-delivered CRISPR/Cas9 gene editing efficiently induced deletion of *MECP2* in the hippocampus of adolescent rhesus monkeys, which elicited core ASD-like phenotypes, including defects in social communication and interaction, hyposensitivity to sensory inputs, hypo-daytime activity, abnormal sleep patterns, abnormal hand motions and social reward deficits. Thus, these results demonstrate that viral-based gene editing in a specific brain region can be applied to rapidly develop monkey models that mimic ASD-like symptoms and shorten the modelling cycle compared with traditional embryonic transgenic or gene editing strategies [9, 26].

This work is consistent with earlier studies demonstrating that deletion of *MECP2* in postmitotic neurons in mice produces deficits in synaptic plasticity and RTT-like behaviors [51]. Furthermore, we extended previous findings by revealing the essential role of hippocampal *MECP2* in the regulation of social behaviors. Intriguingly, deletion of *MECP2* in the hippocampus of the monkeys was not sufficient to recapitulate stereotypic behavior, which is observed in *MECP2* transgenic and mutant monkeys [9, 26]. These results suggest that different brain areas have different functions in ASDs and likely mediate distinct ASD symptoms. For example, specific knockdown of *MECP2* in the basolateral amygdala can produce heightened anxiety-like behaviors rather than abnormalities in social interaction [52]. This indicates that autistic-like symptoms could be studied separately in different brain regions using our model, thus shedding light on the connections between genes and specific neural circuitries.

Restoration of *Mecp2* during adulthood has been shown to rescue behavioral and physiological defects in *Mecp2* knockout (KO) or overexpression (OE) mice [53–58]. The reason why defects caused by loss of *MECP2* in the brain are reversible after birth may be due to the fact that MeCP2 regulates brain development through transcriptional repression, as well as regulation of microRNA processing. Restoration of *Mecp2* to the normal level may be able to restore the impaired synaptic plasticity in particular neural circuits by correction of abnormal gene expression caused by *loss*- or *gain-of-function* of *Mecp2* in mouse models. The present findings suggest that the neural circuitry responsible for social behaviors in non-human primates is also plastic and could be manipulated during adolescence. The feasibility of postnatal social-circuit manipulation in primates suggests a possible way to reverse autistic defects in human RTT and ASD patients.

## Materials and Methods

Complete and detailed experimental procedures are provided in the supplementary materials

### Experimental Design

This study aimed to improve the feasibility of inducing ASD models in wild-type monkeys in order to fulfil the large production demands of disease and drug screening studies. Thus, we induced genetic mutations of the ASD-related gene *MECP2* in the hippocampus of adult rhesus macaques *in vivo* via adeno-associated virus (AAV9)-delivered *Staphylococcus aureus* Cas9. Through behavioral analysis, our model successfully mimicked the symptoms of ASD disease, thus paving the way for the rapid generation of genetically engineered non-human primate models for neurobiological studies and therapeutic development.

### Animals and ethics

Eleven young healthy male rhesus macaques (*Macaca mulatta*) (3–5 kg, 2–3 years old) from the Kunming Primate Research Center of the Chinese Academy of Sciences were used in this study. All monkeys were treated in accordance with the National Institute of Health’s (USA) Guide for the Care and Use of Laboratory Animals [59]. All animals had continuous access to water and monkey chow supplemented with seasonal fresh fruit and/or vegetables. The animals lived in their respective social groups (6–8 male monkeys of similar age) for at least five months prior to the experiments. They were transferred to individual cages (0.74 × 0.71 × 0.74 m) after operation for recovery and individual behavioral assessment under normal conditions (temperature 21 ± 2 ºC, 14/10 h light/dark cycle, light from 07:00 am to 19:00 pm, relative humidity 60%, with standard monkey chow and daily fruit). All experimental protocols and animal care and handling were approved by the Ethics Committee of the Kunming Institute of Zoology and Kunming Primate Research Center (NO. IACUC18019), Chinese Academy of Sciences (AAALAC accredited).

### MRI-guided hippocampus localization

The MRI-guided hippocampus localization procedure was adapted from our previous study [29]. Briefly, two rigid glass tubes (3–4 cm long, internal diameter 0.5 mm, external diameter 0.9 mm) were filled with glycerol to serve as markers, appearing as a high signal in the MRI (T2) images.

The markers were fixed onto the skull by dental cement according to stereotaxic coordinates: i.e., anterior-posterior +10 mm and medial-lateral ±10 mm. After operation, the monkeys were sent for MRI scanning [(1) Siemens Avanto 1.5T scanner (Siemens, Erlangen, Germany), 3D FLASH (fast low-angle shot) spoiled gradient-echo sequence, slice 1.0 mm, distance factor 0.2 mm/20%, FOV 150 × 122 mm, resolution 320 × 320, TR 13.0 ms, TE 5.07 ms, average 3. (2) UIH uMR770 3T scanner (United Imaging, Shanghai, China), 3D five-echo GRE sequence, slice = 0.8 mm, distance factor 0.1 mm/10%, FOV = 124 × 124 mm, resolution 320 × 320, TR = 2300 ms, TE = 380 ms, average 2]. The size of the hippocampus and its relative location to the glycerol markers were calculated using the MRI images (Fig. S1). The injection pathways and volumes were then determined according to the injection coordinates for each monkey listed in Table S2.

### Viral vector design and AAV production

The AAV-SaCas9 vector was obtained from Addgene (RRID: Addgene_61593). The TBG promoter was replaced with human synapsin promoter for better expression in the brain. The sgRNAs were annealed from DNA oligos with BsaI digest ends. After annealing, the sgRNA segments were inserted into the BsaI site of the vector. The AAV9 (AAV-hSyn-SaCas9-U6-sgMECP2-a titer = 1.12 E + 13 V.G./mL; AAV-hSyn-SaCas9-U6-sgMECP2-b titer = 1.64 E + 13 V.G./mL; AAV-hSyn-SaCas9-U6-sgLacZ titer = 1.56 E + 13 V.G./mL; AAV-hSyn-GFP titer = 2.18E + 13 V.G./mL) was constructed at the ION Virus Production Core Facility (Shanghai, China).

### Statistical analysis

Data analysis was conducted using GraphPad Prism v8.00 (GraphPad Software, La Jolla, California, USA), SPSS v19.0 (SPSS, Chicago, IL, USA) and R (version 3.3.2) for Windows. Mann Whitney tests were applied for intragroup analysis. Repeated-measures analysis of variance (ANOVA) was used for intergroup analysis. For modified brinkman board test, the effect of group, size and size-by-group interactions of brinkman score were assessed using a general linear model (GLM) with the following equation: dependent variable (brinkman) = Intercept +group + size + group×size+ random error. The group effect p values by corrections for multiple tests were conducted using Bonferroni. Data are means ± SEM, * *p* < 0.05, ** *p* < 0.01, *** *p* < 0.001.

## Acknowledgments

We thank the members of the Neuroscience Pioneer Club for valuable discussions. We thank the ION Virus Core Facility for producing the AAV virus.

## Author contributions

Zi-Long Qiu and Xin-Tian Hu designed and led the project. Shi-Hao Wu performed virus injection surgery. Xiao Li and Tian-Lin Cheng designed the Cas9 system and tested the virus. Shi-Hao Wu, Dong-Dong Qin, and Lin-Heng Zhang performed behavioral experiment. Zhi-Fang Chen and Shi-Hao Wu performed histological tests. Zi-Long Qiu, Shi-Hao Wu, and Xin-Tian Hu prepared the manuscript.

## Supplementary Materials

1.Supplemental Experimental Procedures

2.Figures S1 to S9

3.Tables S1 to S3

4.Movie details

5.Supplemental References

## Experimental procedures

### Animals and ethics

Eleven young healthy male rhesus macaques (*Macaca mulatta*) (3–5 kg, 2–3 years old) from the Kunming Primate Research Center of the Chinese Academy of Sciences were used for this study. All monkeys were treated in accordance with the National Institute of Health’s (USA) Guide for the Care and Use of Laboratory Animals [1]. All animals had continuous access to water and monkey chow supplemented with seasonal fresh fruit and/or vegetables. All animals lived in their respective social groups (6–8 male monkeys of similar age) for at least five months prior to the experiments. They were transferred to individual cages (0.74 × 0.71 × 0.74 m) after operation for recovery and individual behavioral assessment under normal conditions (temperature 21 ± 2 ºC, 14/10 h light/dark cycle, light from 07:00 am to 19:00 pm, relative humidity 60%, with standard monkey chow and daily fruit). All experimental protocols and animal care and handling were approved by the Ethics Committee of the Kunming Institute of Zoology and Kunming Primate Research Center (NO. IACUC18019), Chinese Academy of Sciences (AAALAC accredited).

### MRI-guided hippocampal localization

The MRI-guided hippocampal localization procedure was adapted from our previous study [2]. Briefly, two rigid glass tubes (3–4 cm long, internal diameter 0.5 mm, external diameter 0.9 mm) were filled with glycerol to serve as markers, appearing as a high signal in the MRI (T2) images. The markers were fixed onto the skull with dental cement according to stereotaxic coordinates: i.e., anterior-posterior +10 mm and medial-lateral ±10 mm. After operation, the monkeys were sent for MRI scanning. [(1) Siemens Avanto 1.5T scanner (Siemens, Erlangen, Germany), 3D FLASH (fast low-angle shot) spoiled gradient-echo sequence, slice 1.0 mm, distance factor 0.2 mm/20%, FOV 150 × 122 mm, resolution 320 × 320, TR 13.0 ms, TE 5.07 ms, average 3. (2) UIH uMR770 3T scanner (United Imaging, Shanghai, China), 3D five-echo GRE sequence, slice = 0.8 mm, distance factor 0.1 mm/10%, FOV = 124 × 124 mm, resolution 320 × 320, TR = 2300 ms, TE = 380 ms, average 2]. The size of the hippocampus and its relative location to the glycerol markers were calculated using the MRI images (Fig. S1). The injection pathways and virus volumes were then determined according to the injection coordinates for each monkey listed in Table S2.

### Viral vector design and AAV production

The AAV-SaCas9 vector was obtained from Addgene (#61593). The TBG promoter was replaced with human synapsin promoter for better expression in the brain. The sgRNAs were annealed from DNA oligos with BsaI digest ends. After annealing, the sgRNA segments were inserted into the BsaI site of the vector. The AAV viruses (AAV-hSyn-SaCas9-U6-sgMECP2-a titer = 1.12 E + 13 V.G./mL; AAV-hSyn-SaCas9-U6-sgMECP2-a titer = 1.64 E + 13 V.G./mL; AAV-hSyn-GFP titer = 2.18E + 13 V.G./mL) were constructed at the ION Virus Production Core Facility (Shanghai, China).

### AAV injections

Prior to surgery, all monkeys were anesthetized with atropine (SFDA approval number: H41021257, 0.03–0.05 mg/kg, i.m.), and then injected with ketamine (Veterinary Drug approval number (2015)100761663, 10 mg/kg, i.m.) and pentobarbital (Merck, 40 mg/kg, i.m.). All surgical procedures were conducted under strict aseptic conditions. As a prophylactic antibiotic treatment, cephalosporin (SFDA approval number: H23021439) was injected for five consecutive days after surgery (25 mg/kg/day, i.m, once a day). Each monkey’s head was fixed on an RWD stereotaxic frame (Product Model: 68902, Shenzhen, China) and a craniotomy was carried out by dental drilling according to the stereotaxic coordinates obtained from MRI scanning. The virus was then infused through a 31-gauge Hamilton syringe placed in a syringe pump (WPI Apparatus, Sarasota, USA) at a speed of 800 nL/min. After infusion, the needle was left in place for 8 min before being slowly retracted from the brain. The same operation was repeated until all injections were finished. After injection, the wound was cleaned thoroughly and sutured [3].

### Immunohistochemistry

Monkeys were anesthetized with pentobarbital (45 mg/kg, i.m) and transcardially perfused with 2000 mL of 4 °C phosphate-buffered saline (PBS) and 500 mL of 4% paraformaldehyde (Sigma-Aldrich, Cat# 16005) in PBS. After perfusion, the hemispheres of the brain were dissected, cut into small blocks, fixed with 4% paraformaldehyde in PBS, and equilibrated in 30% sucrose. The fixed and equilibrated brain tissue blocks were then cut into 50-μm cortical sections with a Microm HM525 cryostat (Microm, Walldorf, Germany). Sections were washed for 5 min in PBS containing 5% bovine serum albumin (BSA) and 0.3% Triton X-100, and incubated with primary antibodies (in PBS with 1% BSA and 0.3% Triton X-100) overnight at 4 °C and subsequently with corresponding secondary antibodies (Alexa-Fluor-conjugated, Invitrogen, at 1:1000). DAPI was used to label the nuclei and sections were mounted with 75% glycerol. Other antibodies used included: GFP antibody (Abcam, ab6673), MeCP2 antibody (Cell Signaling, 3456S), anti-GFAP (1:800; Biolegend, SMI-21R), and anti-Iba1 (1:800; Wako, 019-19741). Confocal imaging and cell quantification were performed. Confocal z-stack images were acquired on a Nikon A1 confocal laser microscope system (Japan). NIH ImageJ software was used to count cell numbers, analyze mean fluorescence intensity of immunoreactive cells, and quantify Iba1 morphology according to previous protocols [4]. Cell counts were performed in nine randomly selected sections in each group.

### Activity and sleep pattern measurement

Actical physical activity monitors (Respironics, Pennsylvania, USA) were used to monitor activity levels of freely moving monkeys during the day (07:00 am–19:00 pm) and sleep patterns during the night (19:00 pm–07:00 am). The monitors are water resistant (IEC Standard 60529 IPX7), lightweight (16 g), and small in size (2.9 × 3.7 × 1.1 cm ^3^). The device was calibrated with monkey information using Actical software before being placed on the monkey’s neck by a soft belt. The monitor was mounted on the same location on each subject to ensure consistency of the data. The monitor uses a single internal omnidirectional accelerometer to sense motion in all directions and integrates the amplitude and frequency of detected motion. The activity data were collected at a sampling rate of 32 HZ and in epochs as short as 1 s. Each monkey was monitored for a period of 7–9 d (according to battery life).

The 7 d of activity was analyzed off-line, and sleep patterns were categorized in 1-min epochs using the night-time data into three states, i.e., awake, transitional, and relaxed sleep [5]. According to previous literature, the animal was considered awake when locomotion occurred or when three or more movements occurred within 1 min. Sleep was scored as transitional when one or two movements occurred within 1 min. The relaxed sleep state was assigned when the animals exhibited no body or limb movements. Sleep duration was the sum of all 1-min epochs of different states (transitional or relaxed sleep) and sleep fragmentation was defined as the frequency of waking bouts lasting 1 min or longer per night.

### Social interaction test

Social interaction tests were conducted, in which behaviors were video-recorded in social groupings (6–8 male monkeys of similar age) and analyzed off-line to assess social interaction changes among different manipulations [5]. In this study, measured behaviors included active social contact, aggressive, submissive, and stereotypical behaviors. Active social contact was defined as initiating social contact, such as playing, grooming, sharing toys, and sitting together (within another monkey’s reach or in contact) between two and/or more monkeys [6]. Aggressive behaviors included biting, grabbing, slapping, open-mouth threats, stare-threats, chasing, and forced displacement. Submissive behavior was defined as screaming, crouching, fleeing, lip smacking, submissive presenting, grimacing, or escaping[7, 8]. Stereotypical behavior was defined as repetitive and consistent actions with no apparent purpose, including digit (finger or toe) sucking, pacing (repetitive, ritualized movement usually involving circling the cage), self-grasping (grabbing or holding onto part of their own body), bouncing (jumping up and down on all four legs), rocking (back and forth movement of the upper body with still feet), body spasms (a quick shake of the body), cage shaking (any vigorous shaking of the cage), and lip-smacking (pursing the lips together and moving them to produce a smacking sound) [9]. Each monkey’s behavior was video-recorded at the same time in the morning (09:30 am–11:30 am) or in the afternoon (14:00 pm–16:00 pm) for 7 d (avoiding rainy days). The behaviors were scored as frequencies and durations per hour by three double-blind viewers. The behaviors were marked when the monkey was unobscured and the viewers could clearly observe behaviors.

### Social reward test

Social reward was studied using the social conditioned place preference (CPP) test, as employed in previous studies [10]. The apparatus for the social CPP test was divided into two equally sized acrylic cages (100 × 100 × 100 cm), with an opening (25 × 40 cm) at the base. The two cages were different colors (white versus red) and could be easily discriminated by the monkey. After 5 d of adaptation to the test chamber (30 min per day per monkey), the formal experiment was commenced. We initiated 30 min of video analysis per day (pre-conditioning trial) to establish color preference (baseline) for the two chambers (white or red). After 3 d of testing, all six monkeys exhibited preference for the red chamber. The monkeys then underwent isolation conditioning for 24 h in the preferred (red) cage, followed by 24 h of social conditioning (with one female cage-mate) three times in the non-preferred (white) cage. After this, each monkey underwent a 30-min postconditioning trial to measure its preference for the two cages. The amount of time spent freely exploring each cage during the 30-min postconditioning trial was recorded. The duration time (time spent) in the social cage was compared between the pre- and post-conditioning trials to demonstrate whether the monkey established preference for the social conditioning cage (Fig. S5).

### Active avoidance test

A modified version of the active avoidance apparatus was used to evaluate fear responses of the monkeys to loud noise. The equipment included two plexiglass observation chambers (80 × 80 × 80 cm) connected by a tunnel (60 × 40 × 20 cm). In each chamber, one remote -controlled high-pitch loudspeaker (116 dB) was installed. A video camera was used to record monkey behavior. Each monkey was tested once daily at the same time for 3 d. Before the test, each monkey was trained to follow a monkey guide stick to enter the test chamber and adapt to free shuttling between the two chambers over several days. Afterwards, the monkey was guided with the stick from the home cage into the test chamber. Each monkey was given 10 min to adapt to the chamber, after which a loud noise (116 dB) was presented, which should elicit an escape response from the monkey to move to the adjoining chamber. The noise was stopped when the monkey escaped to the other chamber. To ensure no harm was inflicted, the noise was stopped after 2 min if the monkey had not successfully escaped. Escape time from the noise chamber to the quiet chamber was recorded [5].

### Gaze-following test

A monkey’s ability to follow another’s gaze can be assessed using a gaze-following test. Unlike adults, juveniles are not able to orient their attention on the basis of another’s gaze alone[11]. Therefore, head-eye cues were presented to the monkeys to determine differences in gaze-following ability between the Cas9-MECP2 and Cas9-Control monkeys. During the test, the experimenter sat facing the monkey at a 1.5-m distance, and then turned his/her head 70°up, down, left, or right and maintained each posture for 3 s, with the eyes always aligned with the head. In the test, for each head orientation, a total of 25 trials were carried out randomly (100 trials per day). Each monkey was tested under freely moving conditions at the same time of the day. The responses of the monkeys were recorded by a video camera positioned behind the experimenter. A trial was considered valid only if the monkey was looking at the experimenter at the beginning of the trial. Each monkey was tested for three consecutive days and the percentage of gaze-following responses (number of responses divided by total number of valid trials in each test session) was calculated. To control for the possible effect of differences in attention to the experimenter between the Cas9-MECP2 and Cas9-Control monkeys, 5 min of behavioral analysis was scored randomly during each test session. The amount of time in which the monkey’s body was oriented toward the experimenter and the monkey visually explored the experimenter was totaled [11] (Fig. S6).

### Modified Brinkman board test

This test, modified and adapted from previous reports, is widely used for assessing manual dexterity in non-human primates [12]. Behavioral assessment of manual dexterity is based on manual grasping tasks (use of precision grip to obtain peanut kernel). Briefly, monkeys retrieve a peanut kernel from 21 slots of different size (four 1 × 2 cm, four 1.5 × 2 cm, four 2 × 2 cm, three 1 × 3 cm, three 1.5 × 3 cm, and three 2 × 3 cm). Two parameters (accuracy and contact time) were analyzed by replaying the recorded video sequences. Accuracy: correct rate of retrievals of peanut kernel from slots. Contact time: time interval between first finger touch of the pellet and onset of retrieval of the pellet from the well. Time interval was measured by replaying the video sequence frame-by-frame.

### Gait speed

Gait speed is a reflection of motor processes[13]. To observe whether the Cas9-MECP2 monkeys exhibited motor disabilities, we measured gait speed following previous study [13]. Briefly, usual walking speed was quantified by offline video analysis from 5 d of interference-free video surveillance recording (09:30 am–16:30 pm). Only monkeys traversing a minimum of 1 m at a normal pace without any motivation, such as food or provocation, were analyzed. Five valid instances per monkey were used for statistical analysis.

### Reversal learning (RL)

Reversal learning tasks are used to assess learning ability and cognitive flexibility[14]. Here, testing was conducted in a Wisconsin General Test Apparatus (WGTA). Monkeys were familiar with the WGTA and fasted before the experiment. First, monkeys were taught to flip the cover and obtain peanuts from the food well. Second, two different shaped covers (“◊” and “☆”) were used to cover two food wells, with the peanut always placed in the food well covered by “☆”. The “☆” was placed (right or left) randomly, and monkeys could only select one of the two shapes (“◊” or “☆”). Only when the monkey selected “☆” was it able to obtain the peanut. When the monkey’s peanut success rate was above 85% in 45 successive trails, we added two new shapes (“○” and “♡”), with the peanut always placed in the food well covered by “♡”. The four shapes appeared randomly with equal probability in a session (16 trials). When the monkey’s peanut success rate was above 85% in three successive sessions, we started the reverse task, where selecting “○” and “◊” achieved peanuts. The number of trials required until the monkey reached an 85% success rate in a session was recorded.

### Pattern-recognition task

A modified version of the pattern-recognition task was used to evaluate memory ability [15]. Testing was conducted in a WGTA. Monkeys were familiar with the WGTA and fasted before the experiment. Five different shaped covers were used to cover five food wells. The position of the cover was randomly decided by the computer, with peanuts only found under one shaped cover. The monkeys first learned two shapes and gradually increased to learning all five shapes. When a monkey’s peanut success rate was above 90% in 60 successive trails, it was considered to have remembered the reward graphic. After one week, the monkey performed the task again, and the correct choice rate of the reward graphic was counted.

### *In vivo* mutation efficiency detection

To detect whether gene editing occurred in the monkey brain tissue near the injection sites, fluorescent-positive tissue was collected from slices and then digested by protease K. Genome DNA was extracted by phenol-chloroform and precipitated by absolute alcohol. The proximal regions of the sgRNA-targeting sites were amplified by PCR, followed by Sanger sequencing. Results were compared with wild-type control to confirm whether the target gene was edited.

### Off-target detection

To detect the possible presence of sgRNA off-targeting, an online database (http://www.rgenome.net/cas-offinder/) was used to predict potential off-target sites. From the many potential off-target sites, several of the most likely sites were selected (Table S3). These selected sites had to satisfy the following conditions: must be located in the *Macaca mulatta* Mmul_10 reference genome database; the five bases on the sgRNA next to PAM (seed region) must be free from mismatch; total number of mismatched bases must be less than or equal to three; and, bulge size must be less than or equal to one. The proximal regions of these selected potential off-target sites were amplified by PCR, followed by Sanger sequencing. Results were compared with wild-type control to confirm whether real off-target sites existed.

## Supplementary Figures

**Figure S1.**
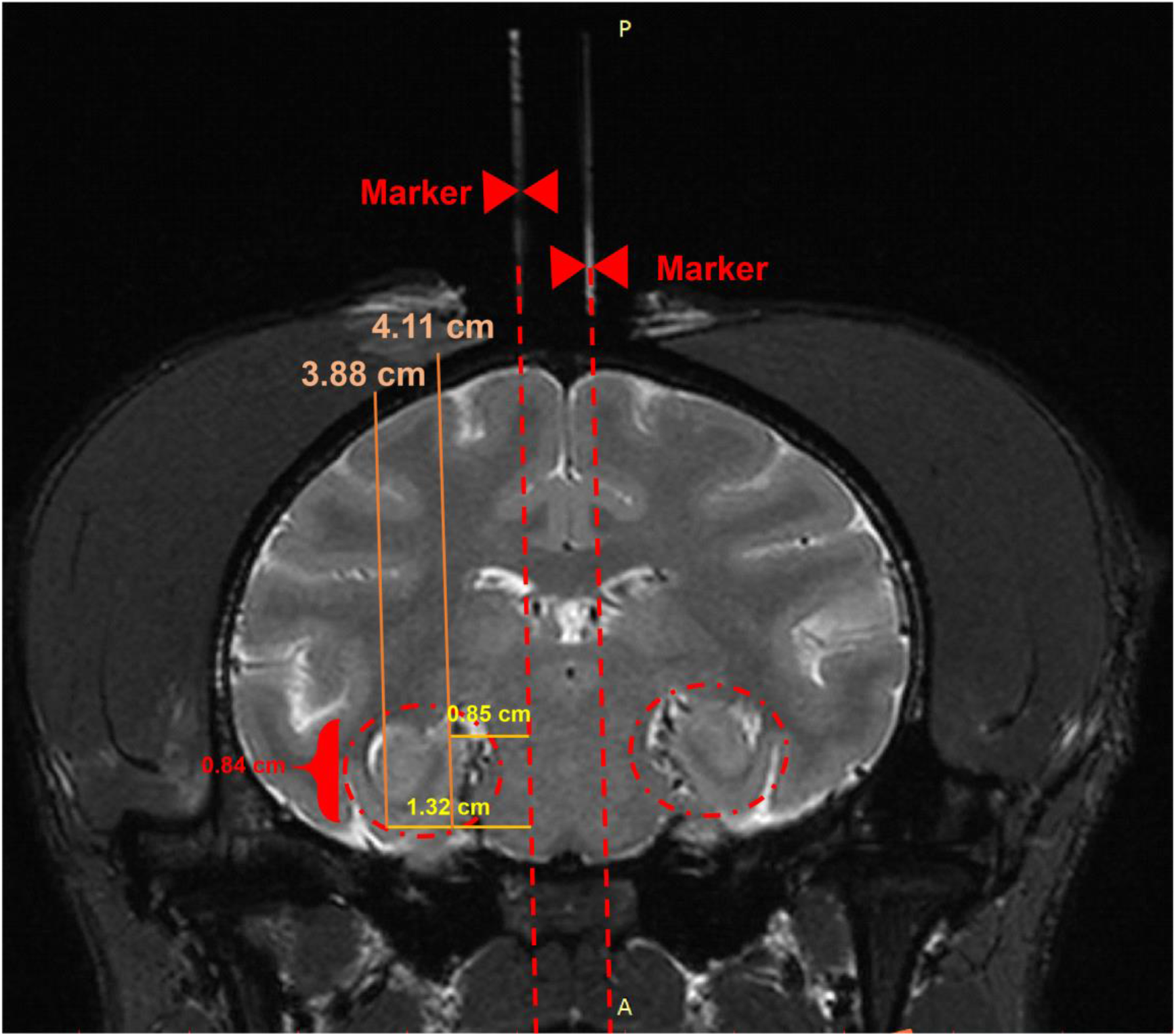
MRI images of marker locations, hippocampus, and coordinates of injection pathways. MRI image displays monkey-skull markers (red arrows), which were used as references to locate hippocampus in coronal plane. Red circles indicate location of hippocampus and red number shows range of injection depths). Orange and yellow numbers show maxima and minima coordinates of injection pathways in this section, respectively.

**Figure S2.**
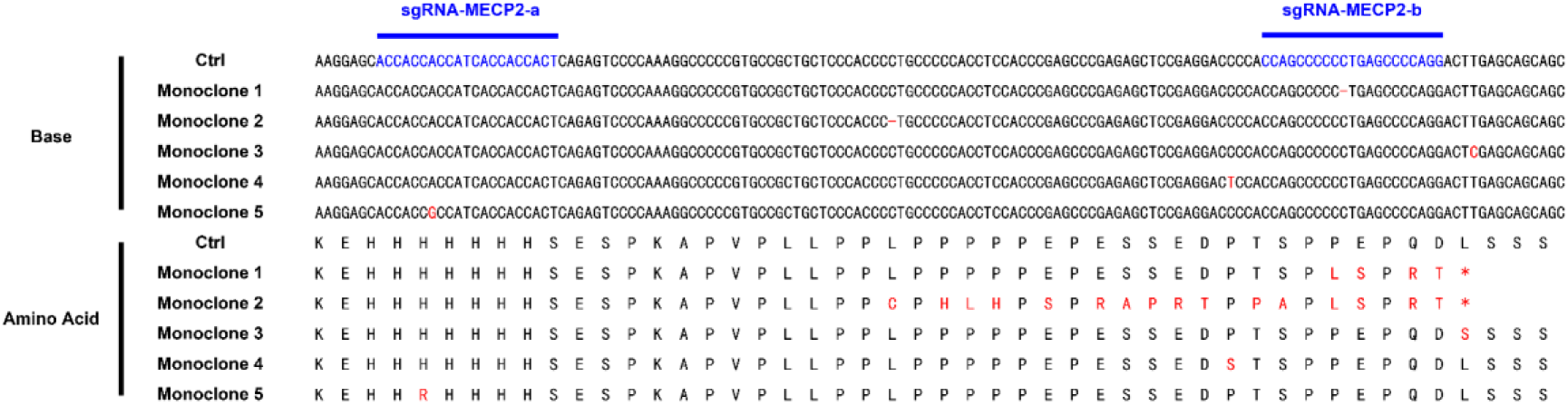
Identification of genetic mutations caused by SaCas9 and sgRNAs. Examples of monoclone mutations near target site caused by sgRNA in Cas9-MECP2 (KO-3) and Cas9-Control monkey (Ctrl-2) brains. Red highlights show mutated base and amino acid, and blue highlights show sgRNA target sites.

**Figure S3.**
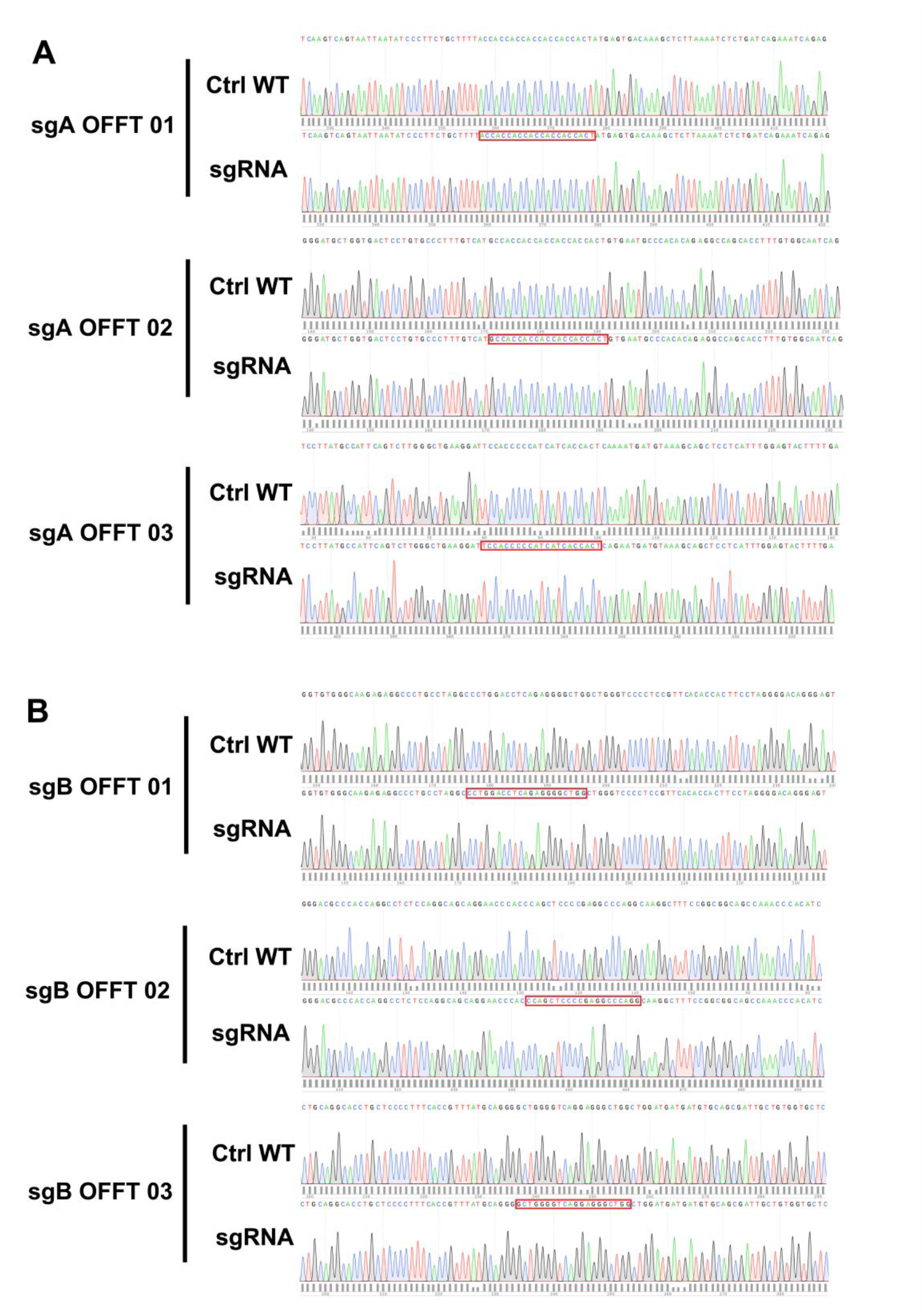
Sequencing results of predicted off-target sites. Potential off-target sites of sgRNA1 (A) and sgRNA2 (B) were predicted using Cas-OFFinder database (http://www.rgenome.net/cas-offinder/). Typical sequencing results are shown and compared with genome sequences of wild-type monkeys.

**Figure S4.**
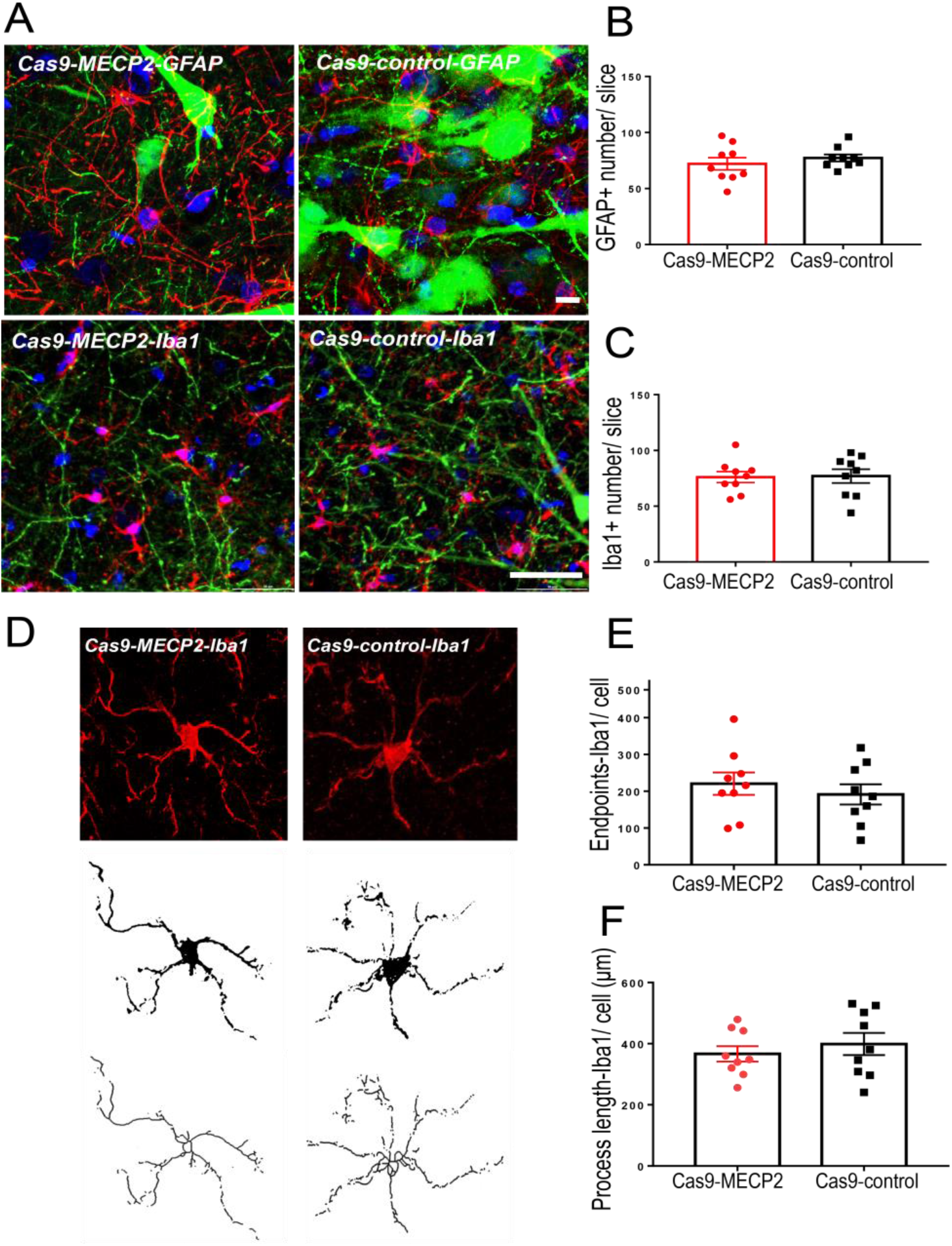
Immunohistochemical analysis of astrocytes and microglial cells. (A) GFAP-positive astrocytes and Iba1-positive microglial cells in AAV-transduced brain area (GFP (green), GFAP (red), Iba1(red), and DAPI (blue); GFAP bar = 10 μm; Iba1 bar = 50 μm). (B) No significant differences were found in number of GFAP-positive astrocytes between Cas9-MECP2 (n = 9 slices) and Cas9-Control groups (n = 9 slices, Mann Whitney test, *p* = 0.4755). (C) No significant differences were found in number of Iba1-positive microglial cells between Cas9-MECP2 (n = 9 slices) and Cas9-Control groups (n = 9 slices, Mann Whitney test, *p* = 0.6196) (D) Example photomicrograph of fluorescent IHC and cropped cell with skeletonized image. There were no significant differences in (E) endpoints (Mann Whitney test, *p* = 0.5302) or (F) process lengths (Mann Whitney test, *p* = 0.5457) in Iba1 cells between Cas9-MECP2 (n = 9) and Cas9-Control groups (n = 9). Data are means ± SEM.

**Figure S5.**
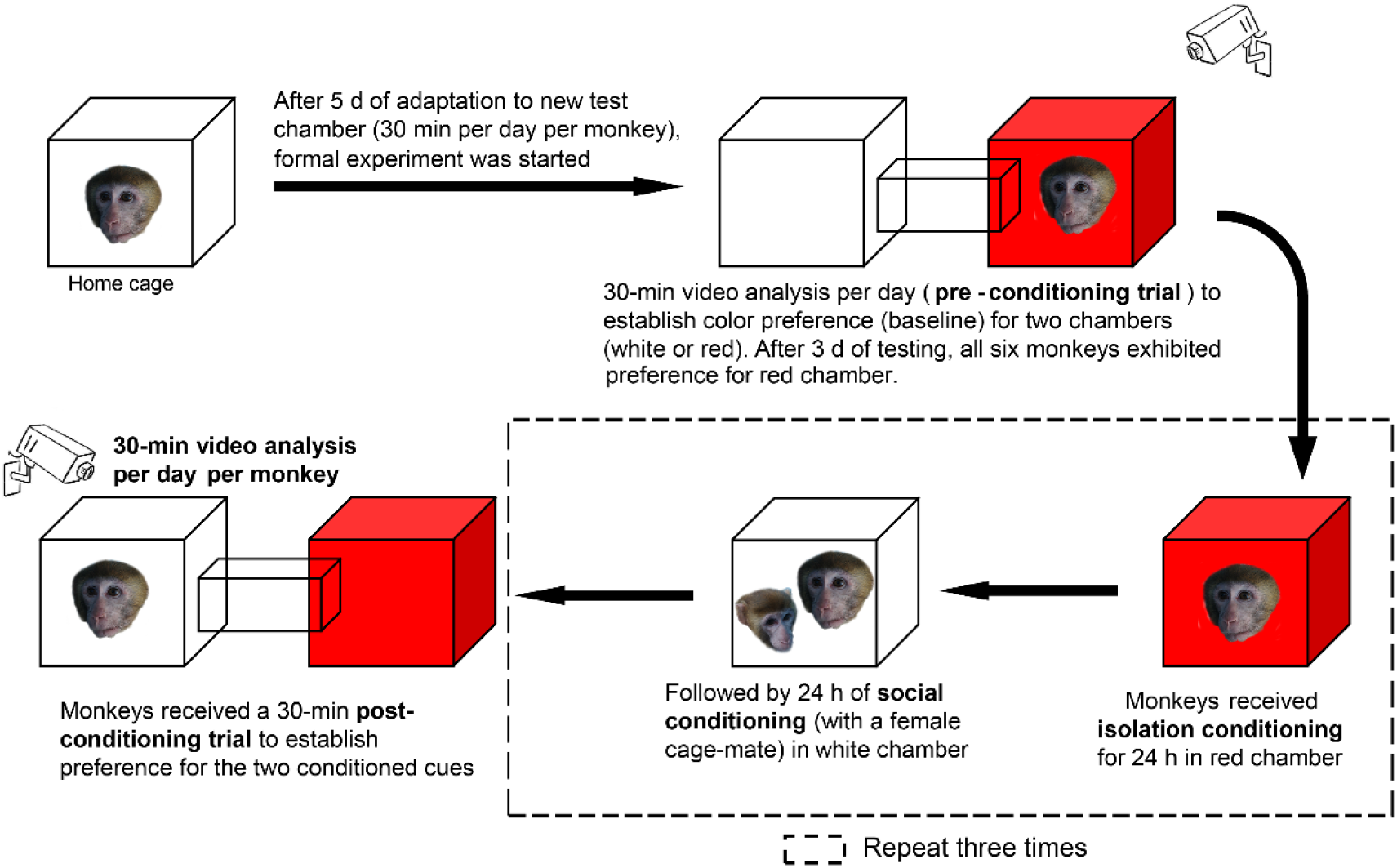
Schematic of social reward test procedure.

**Figure S6.**
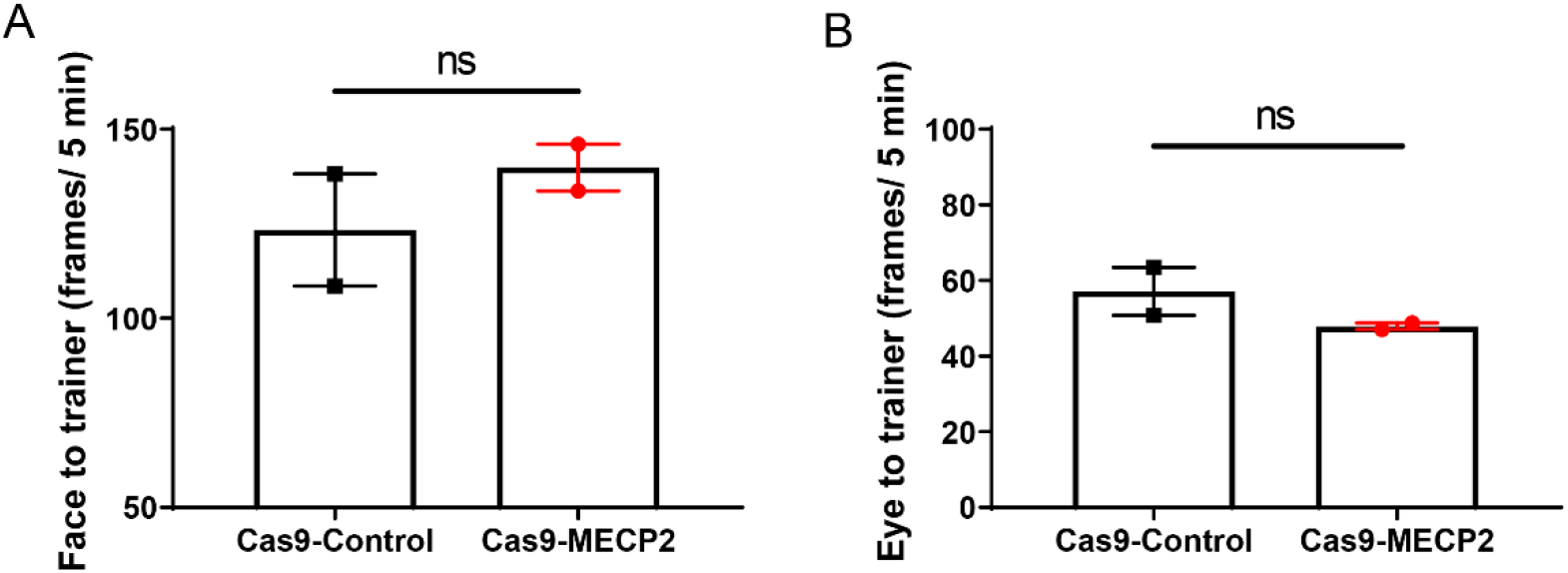
Controls for GFR test. There were no significant differences between Cas9-MECP2 and Cas9-Ctrl monkeys in: (A) amount of time in which monkey’s body was oriented toward experimenter (Mann Whitney test, *p* = 0.6667) and (B) monkey looked at experimenter (Mann Whitney test, *p* = 0.33). Data are means ± SEM.

**Figure S7.**
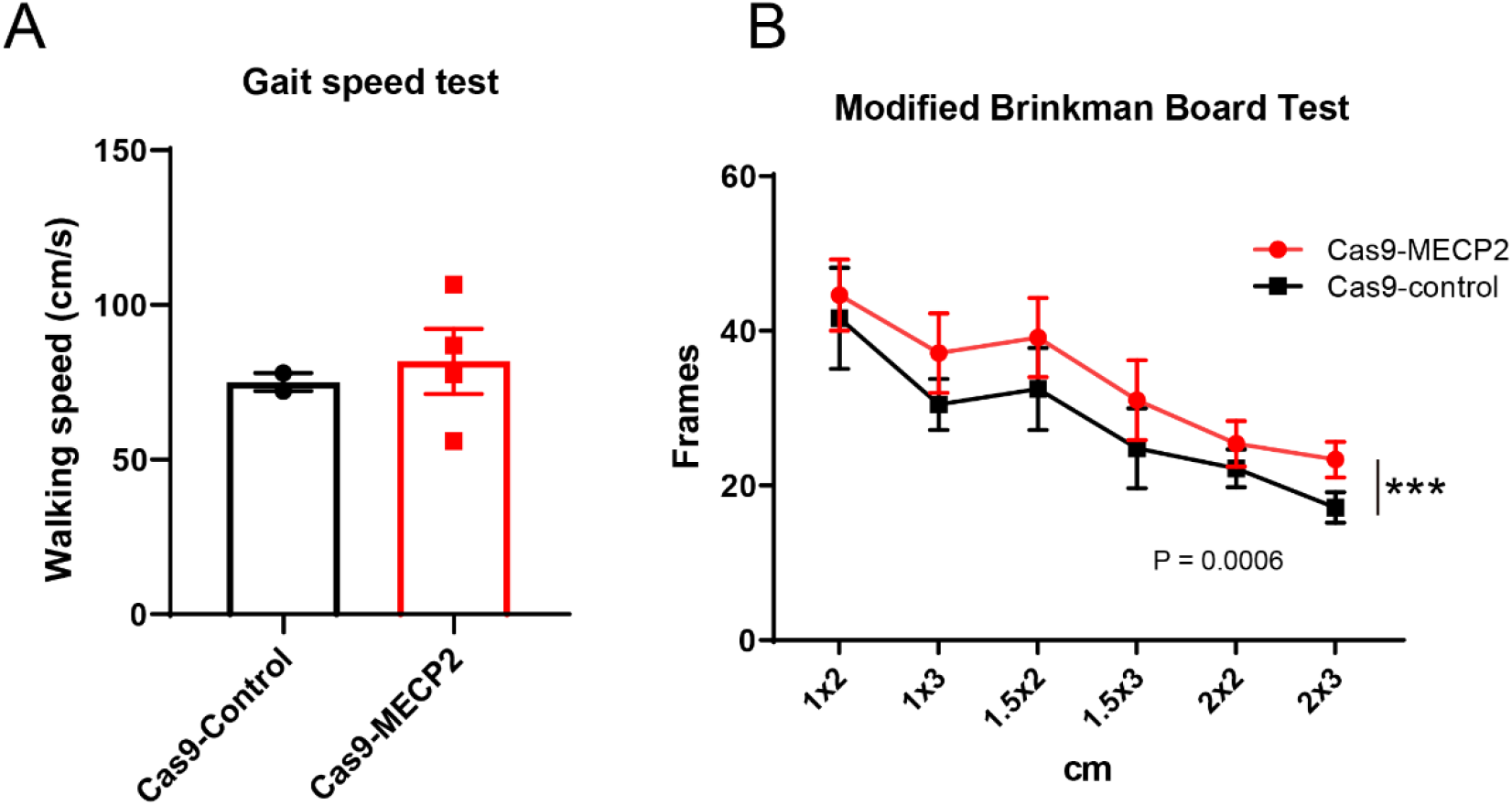
(A) Walking speed (cm/s) of Cas9-MECP2 (n = 4) and Cas9-Control groups (n = 2). Average of five samples over 5 d of walking speed test is depicted (Mann Whitney test, *p* = 0.3675), data are means ± SEM. (B) Modified Brinkman board test was used to assess manual dexterity. There was significant difference in latency to grab peanut kernel from grid between Cas9-MECP2 (n = 5) and Cas9-Control groups (n = 3, *p* = 0.0006). Average of 21 samples per monkey over three consecutive days is depicted, data are means ± SEM..

**Figure S8.**
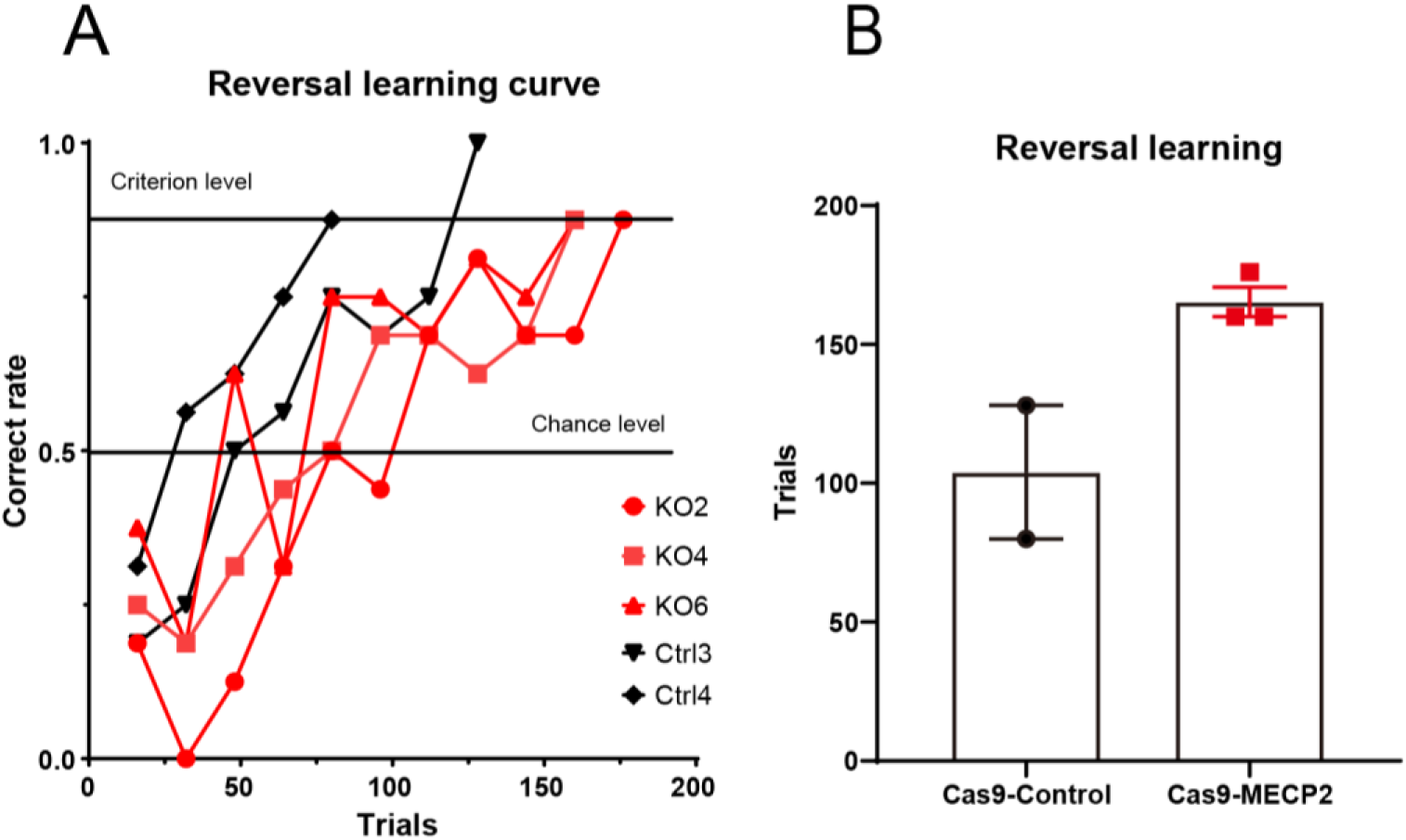
Performance of reversal learning task of two groups of monkeys. (A) Learning curve with reverse task in Cas9-MECP2 (n = 3) and Cas9-Control monkeys (n = 2), and (B) number of final trials to reach criterion level showed no significant differences between Cas9-MECP2 and Cas9-Control groups (*p* = 0.1). Data are means ± SEM. Non-parametric tests (Mann Whitney) were used for statistical analysis.

**Figure S9.**
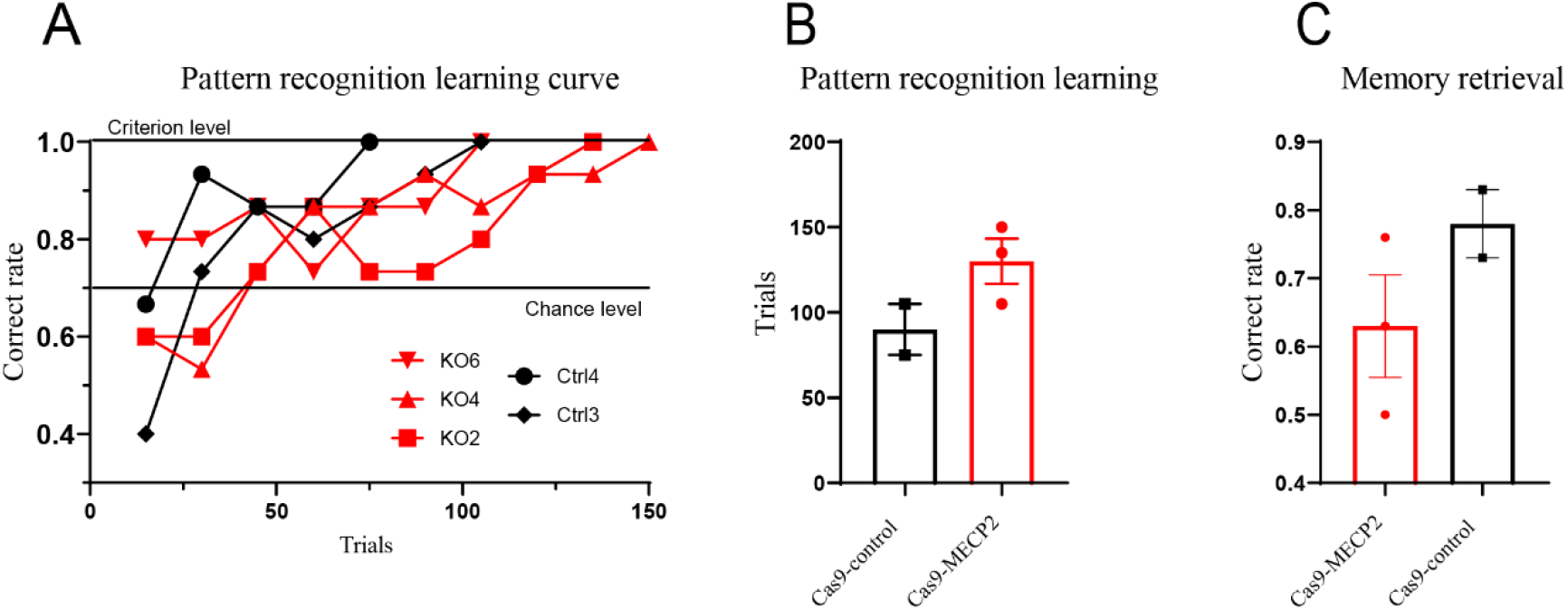
Performance of pattern recognition and memory retrieval tasks of monkeys with gene editing. During learning phase, (A) learning curve and (B) number of final trials to reach criterion level showed no significant differences between Cas9-MECP2 (*p* = 0.3, n = 3) and Cas9-Control groups (n = 2). During memory retrieval phase, (C) there was no significant difference in correct rate between groups (*p* = 0.4). Data are means ± SEM. Non-parametric tests (Mann Whitney) were used for statistical analysis.

**Figure S10.**
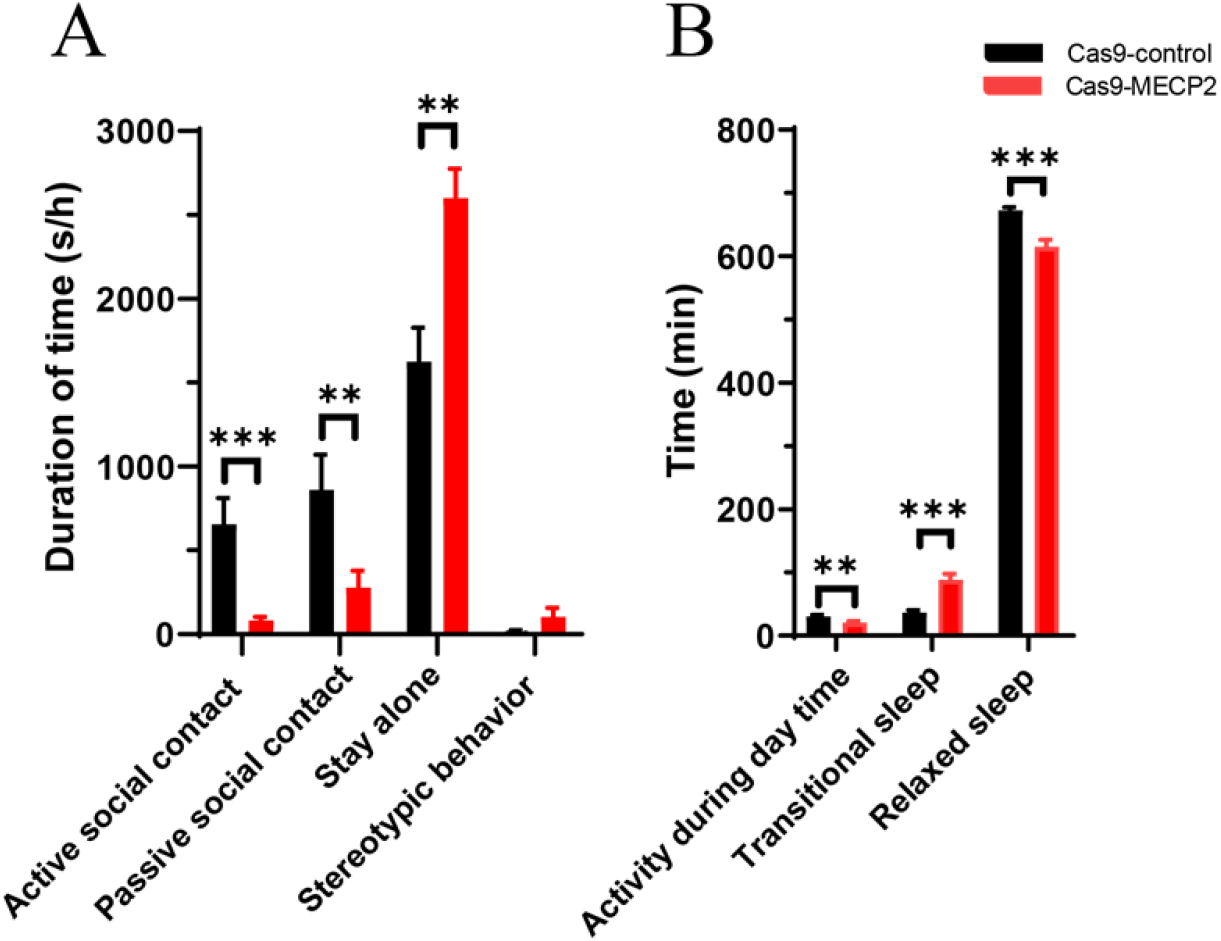
longitudinal analysis of social behaviors, light/dark activities, and sleep patterns consistently showed ASD phenotypes. (A) Duration of active social contact (*p* = 0.0002), passive social contact (*p* = 0.0047), stay alone (*p* = 0.0038), and stereotypic behavior (*p* = 0.143) of Cas9-MECP2 and Cas9-Control monkeys. (B) Compared to Cas9-contorl monkeys, Cas9-MECP2 monkeys showed decreased day-time activities (*p* = 0.0028), longer transitional sleep time (*p* < 0.0001), and shorter relaxed sleep times (*p* < 0.0001). Data were averaged over 5 d and showed means ± SEM, * *p* < 0.05, ** *p* < 0.01, *** *p* < 0.001. Non-parametric test (Mann Whitney) was used for statistical analysis.

**Table S1.** Information on individual monkeys for behavioral tests.

**Table S1A.** Social interaction and stereotypic behavior.

| ID | Age at gene editing (years old) | Days after gene editing |
| --- | --- | --- |
| Ctrl-1 | 3 | 93 |
| Ctrl-2 | 3 | 91 |
| Ctrl-3 | 2 | 109 |
| KO-1 | 3 | 157 |
| KO-2 | 2 | 154 |
| KO-3 | 3 | 97 |
| KO-4 | 3 | 95 |
| KO-5 | 3 | 93 |
| KO-6 | 3 | 92 |
| KO-7 | 3 | 91 |

**Table S1B.** Light activity and sleep patterns.

| ID | Age at gene editing (years old) | Days after gene editing |
| --- | --- | --- |
| Ctrl-1 | 3 | 93 |
| Ctrl-2 | 3 | 91 |
| Ctrl-3 | 2 | 109 |
| KO-3 | 3 | 98 |
| KO-4 | 3 | 95 |
| KO-5 | 3 | 93 |
| KO-6 | 3 | 92 |

**Table S1C.** Social reward test.

| **ID** | **Age at gene editing (years old)** | **Days after gene editing** |
| --- | --- | --- |
| Ctrl-2 | 3 | 507 |
| Ctrl-3 | 2 | 256 |
| Ctrl-4 | 2 | 247 |
| KO-4 | 3 | 633 |
| KO-6 | 3 | 198 |
| KO-7 | 3 | 190 |

**Table S1D.** Gaze following test.

| **ID** | **Age at gene editing (years old)** | **Days after gene editing** |
| --- | --- | --- |
| Ctrl-3 | 2 | 95 |
| Ctrl-4 | 2 | 91 |
| KO-6 | 3 | 101 |
| KO-7 | 3 | 97 |

**Table S1E.** Active avoidance test.

| **ID** | **Age at gene editing (years old)** | **Days after gene editing** |
| --- | --- | --- |
| Ctrl-3 | 2 | 138 |
| Ctrl-4 | 2 | 139 |
| KO-6 | 3 | 108 |
| KO-7 | 3 | 105 |

**Table S1F.**
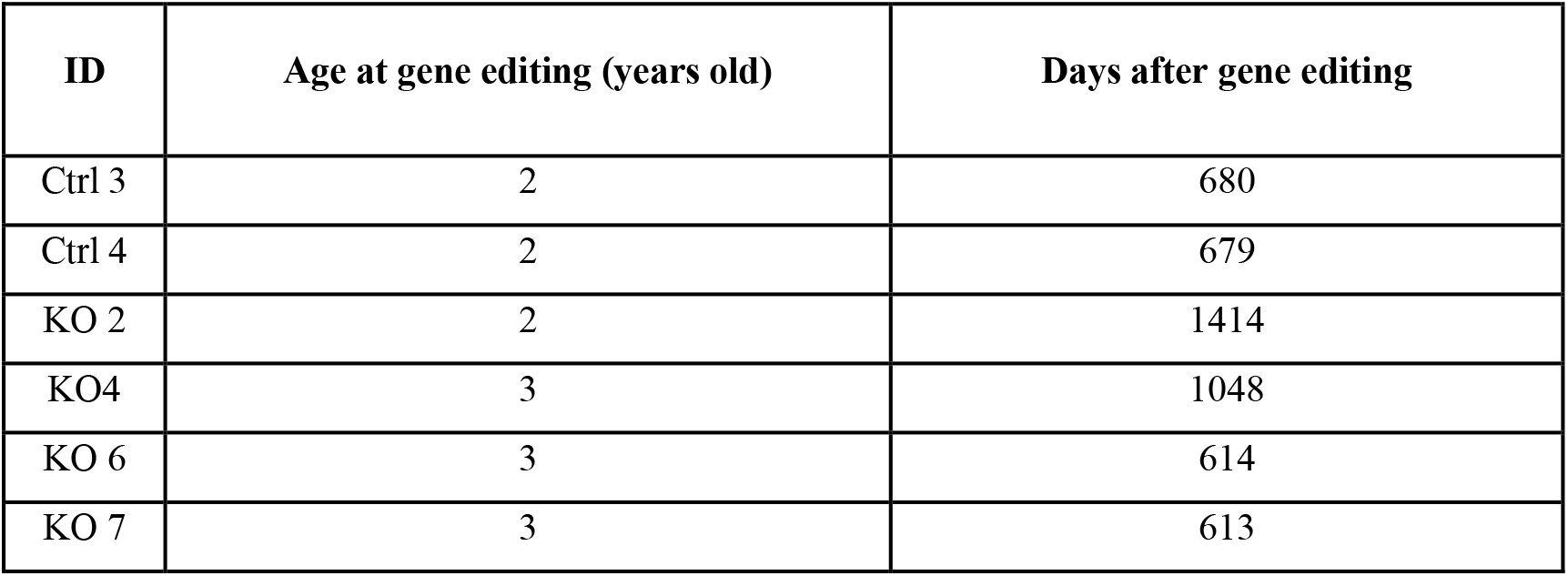
Gait speed test.

**Table S1G.** Brinkman board test.

| **ID** | **Age at gene editing (years old)** | **Days after gene editing** |
| --- | --- | --- |
| Ctrl-2 | 3 | 384 |
| Ctrl-3 | 2 | 124 |
| Ctrl-4 | 2 | 125 |
| KO 2 | 2 | 753 |
| KO4 | 3 | 416 |
| KO 5 | 3 | 298 |
| KO 6 | 3 | 121 |
| KO 7 | 3 | 125 |

**Table S1H.** Reversal learning.

| **ID** | **Age at gene editing (years old)** | **Days after gene editing** |
| --- | --- | --- |
| Ctrl 3 | 2 | 708 |
| Ctrl 4 | 2 | 707 |
| KO 2 | 2 | 1442 |
| KO4 | 3 | 1076 |
| KO 6 | 3 | 642 |

**Table S1I.**
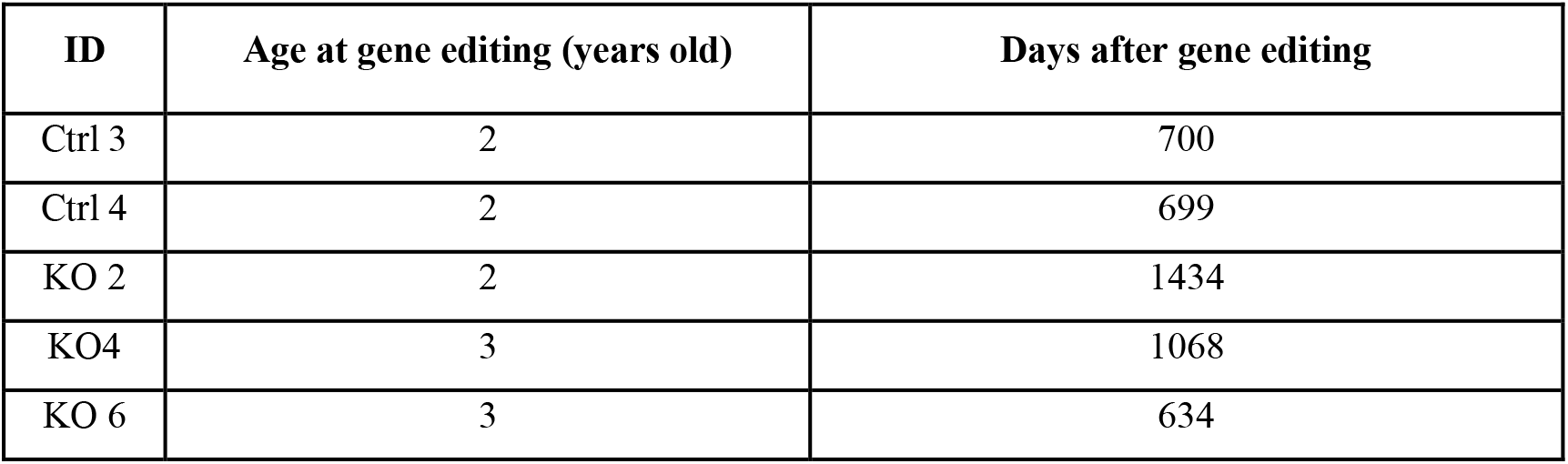
Pattern recognition learning.

| **ID** | **Age at gene editing (years old)** | **Days after gene editing** |
| --- | --- | --- |
| Ctrl 3 | 2 | 700 |
| Ctrl 4 | 2 | 699 |
| KO 2 | 2 | 1434 |
| KO4 | 3 | 1068 |
| KO 6 | 3 | 634 |

**Table S2.** Median value of injection coordinates of each monkey determined from MRI scans.

| Monkey | Coordinates: Median of anterior-posterior/medial-lateral/deep/angle | Total injected vector volume (μL) for bilateral hippocampus |
| --- | --- | --- |
| KO1 | -1.3, -1, -0.9, -0, +0.3 +1 cm / ±12.5, ±16 cm /-2.9 ~ -3.8 cm / 0° | 368 |
| KO2 | -1, -0.7, -0.4, -0.1, +0.2 cm / ±1.36 cm, ±1.80 cm /-2.75 ~ -3.61 cm / 0° | 446 |
| KO3 | -0.7, -0.5, -0.3, -0.15 and +0.1 cm/ ± 1.25, ± 16 cm /-2.9~ -3.7 cm /0° | 386 |
| KO4 | -1, -0.8, -0.6, -0.4, -0.2 / ±1.30, ±1.63 cm /-3.2 ~ -3.8 cm /0° | 392 |
| KO5 | -1.5, -1.3, -1.1, -0.9, -0.7m/ ±1.31, ±1.58 cm /-2.70~-3.49 cm /0° | 372 |
| KO6 | -1.6, -1.2, -0.8, -0.5 cm/ ±1.20, ±1.73 cm /- 3.24~ -3.99 cm / 0° | 398 |
| KO7 | -0.9, -0.7, -0.5, -0.3, -0.1 cm/ ±1.08, ±1.64 cm / -3.31~ - 4.30 cm / 0° | 388 |
| Ctrl | -2.3, -1.9, -1.6, -1.3 cm/ ±1.1, ±1.5 /-3.2~ -4.1 cm /0° | 402 |
| Ctr2 | -0.9, -0.6, -0.3, 0, +2 cm/ ±1.27, ±1.69 /-2.82~ -3.54 cm /0° | 374 |
| Ctr3 | -1.2, -0.9, -0.6, -0.3 cm/ ±1.35, ±1.65 cm / -3.31~ -3.82 cm / 0° | 400 |
| Ctr4 | -0.8, -0.6, -0.4, -0.2, 0 cm/ ±1.13, ±1.49 cm / -2.93~ -3.51 cm / 0° | 388 |

Note: Locations depict: distance before (-) or after (+) interaural (cm); distance left (+) or right (-) to midsagittal plane (cm); distance under (-) skull (mm); and angle from vertical line (°).

**Table S3.**
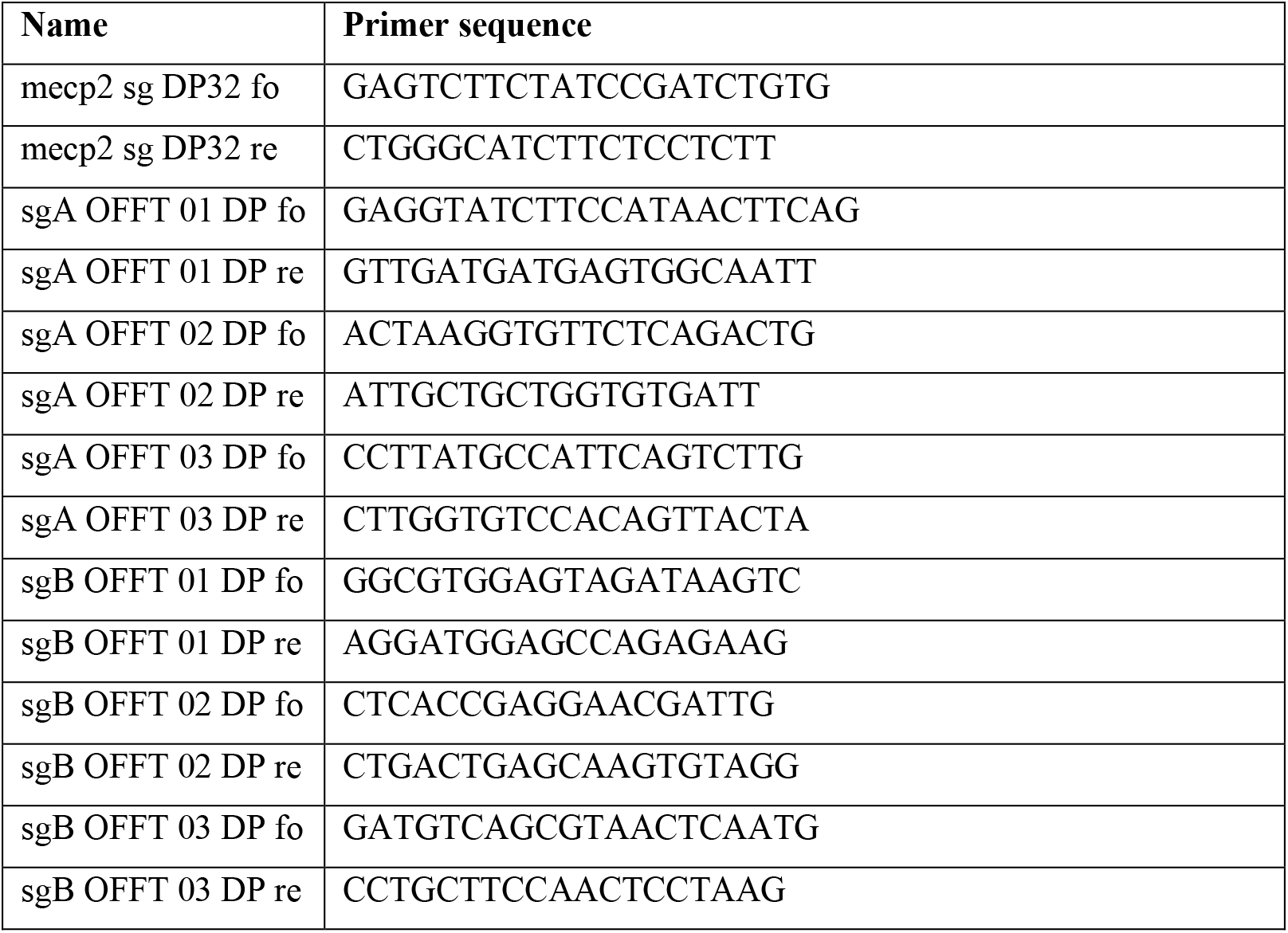
Potential off-target sites predicted by Cas-OFFinder database.

**Table S4.**
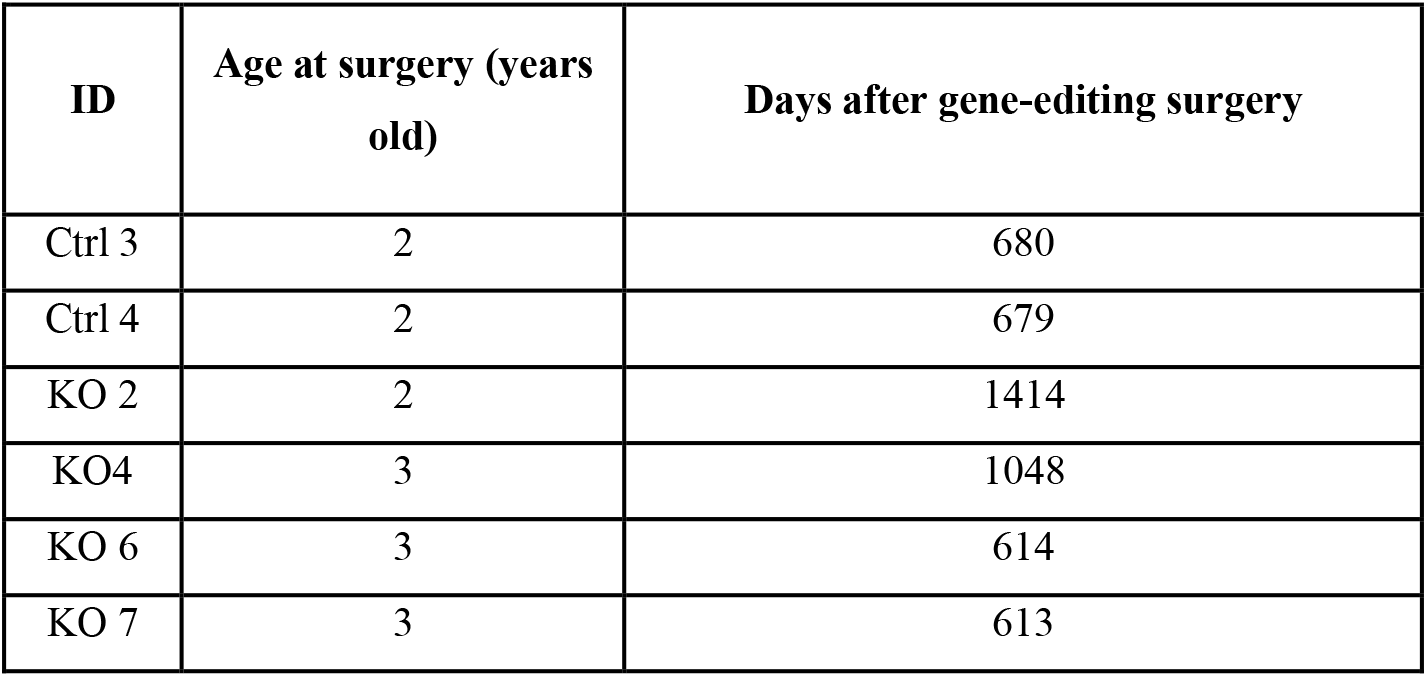
Ages of monkeys during social interaction, light activity, and sleep pattern during revision

| ID | Age at surgery (years old) | Days after gene-editing surgery |
| --- | --- | --- |
| Ctrl 3 | 2 | 680 |
| Ctrl 4 | 2 | 679 |
| KO 2 | 2 | 1414 |
| KO4 | 3 | 1048 |
| KO 6 | 3 | 614 |
| KO 7 | 3 | 613 |

## Movie details

**Movie S1-3. Representative social behaviors changes of gene-edited monkey before and after gene-editing:** macaque showed close social interactions (grooming) with other macaques before virus injection (movie S1). After virus injection, the same macaque spent more time alone in corner (movie S2) and ignored social interactions initiated by other macaques (movie S3)

**Movie S4-5. Representative social reward behaviors:** after social conditioning, control macaque was more willing to stay in white social cage than previously preferred red cage (movie S4), while gene-edited macaque was still willing to stay in previously preferred red cage (movie S5).

**Movie S6-7. Representative gaze-following responses:** control monkey followed direction of experimenter’s gaze (movie S6), while gene-edited monkey did not follow direction of experimenter’s gaze (movie S7).

**Movie S8-9. Representative active avoidance responses:** when alarm sounded, control macaque quickly escaped to another cage (movie S8), while gene-edited macaque showed insensitivity to noise, and took a long time to escape to another cage (movie S9).

